# Dale’s law, stability and nonlinearity are sufficient constraints to ignite transient and self-sustained neural dynamics

**DOI:** 10.64898/2026.09.14.751508

**Authors:** Nicoleta Condruz, Ivan Bulygin, Chaitanya Chintaluri, Tim P. Vogels

## Abstract

The motor cortex orchestrates a rich and flexible repertoire of network dynamics for rhythmic and goal-directed movements. Computational studies have begun to illuminate the mechanistic origins of this repertoire, but a comprehensive model that can explain the emergence of both transient and self-sustained dynamics is still missing. Here, we show that three simple ingredients — Dalean connectivity, stability, and nonlinear neural responses — suffice to reverse-engineer networks that produce transient, steady-state, and self-sustained periodic activity. A single dynamical principle underlies this repertoire: the interaction of non-normal amplification, inherent to Dalean networks, with neuronal nonlinearity, so to ignite and sustain multi-stable, controllable dynamics. Our approach yields entire families of connectivity matrices that require no hand-tuning or learning of weights. Without fitting them to data, these networks reproduce the population-level signatures of motor cortex, implying that the richness of cortical dynamics need not be sculpted by learning, but may emerge from simple biological ingredients.

## Introduction

At the heart of the flexible repertoire of movements that motor cortex can create are distinct patterns of transient and self-sustained neural activity (Fig. 1A).^1–9^ For example, goal-directed arm reaching is driven by complex, multiphasic firing-rate patterns that relax to baseline once the movement is complete.^1,5,10–12^ Rhythmic arm cycling, on the other hand, is sustained by periodic population trajectories.^6,8^ Yet, the computational principles that allow transient and self-sustained activity to emerge in the same network remain elusive.

**Figure 1.**
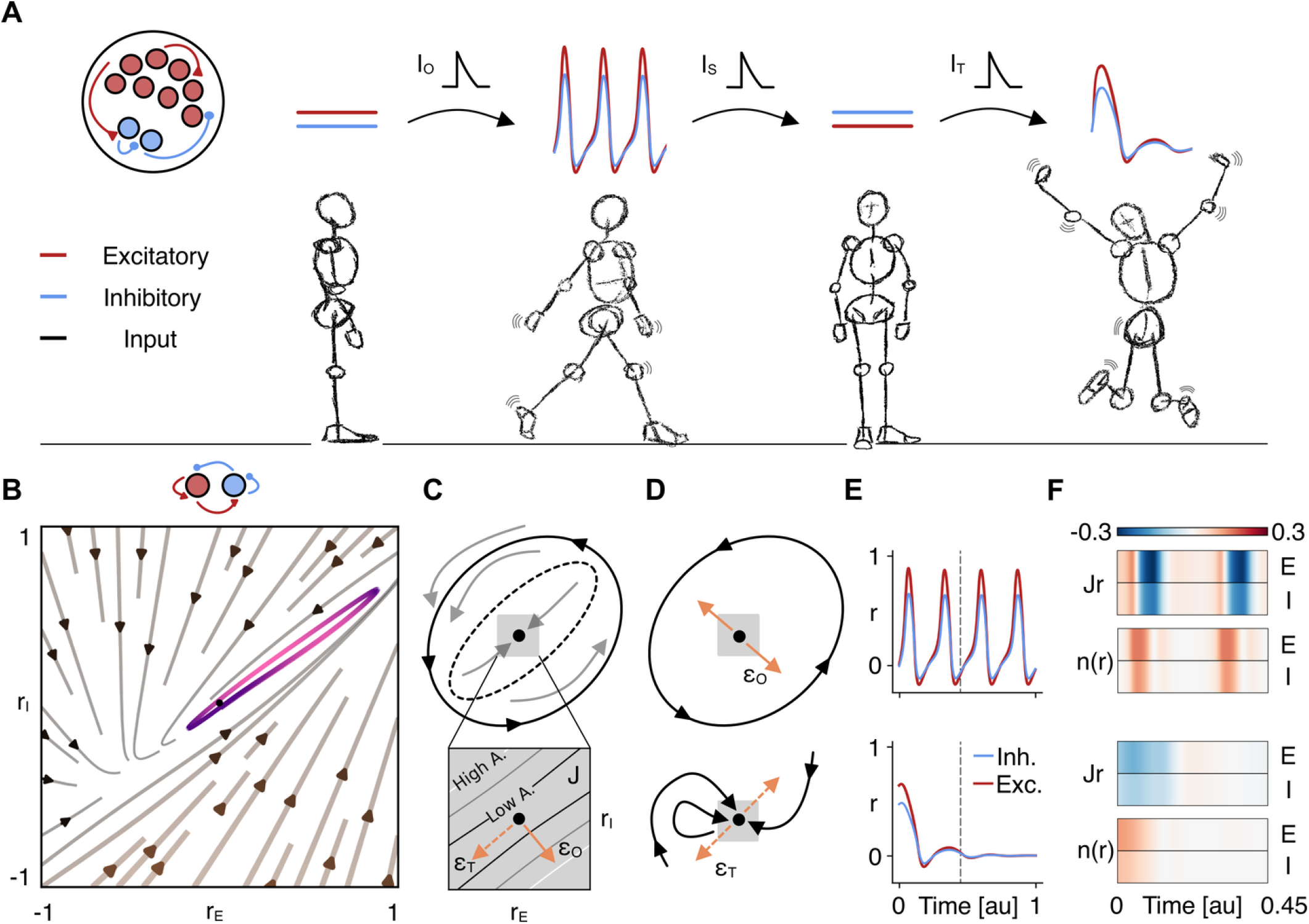
Transient amplification and neuronal nonlinearity sustain persistent neural dynamics. **A)** Illustration of a motor sequence and the corresponding example traces of the neural activity patterns and transition commands necessary to generate it in an E/I network. **B)** Example of a two-dimensional vector field (arrows) spanned by 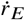 and 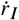 where a stable periodic orbit (purple) surrounds the baseline fixed point (black). **C)** Illustration of the phase-space geometry in which a stable fixed point (center) and a stable limit cycle (solid line) can coexist, separated by an unstable periodic orbit (separatrix, dashed). The gray square denotes the local neighborhood of the fixed point within which the nonlinear dynamics are well approximated by the linearized system. Inset: magnified view of the linear regime showing the effect of the Jacobian *J* of the connectivity. The non-normality of *W* induces a direction-dependent effect where different perturbation directions lead to high amplification (*ϵ*_*O*_) or low amplification (*ϵ*_*T*_ ) of input activity. **D)** Illustration of phase-space trajectories following perturbations aligned with amplifying (*ϵ*_1_, top) and non-amplifying (*ϵ*_2_, bottom) directions. **E)** Neural activity traces showing sustained oscillations following an amplifying perturbation (top), and decay of activity back to baseline following a non-amplifying perturbation (bottom). **F)** Decomposition of amplified (top) and non-amplified (bottom) dynamics into linear (*Jr*) and nonlinear (*n*(*r*)) components (Eq. 2).

Computational models have long sought to uncover these principles through two complementary approaches. Either networks are trained to reproduce cortical activity, or they are built from mechanistic constraints. Trained recurrent networks can host a vast variety of dynamics,^13–22^ but do not necessarily reveal the principles through which they emerge. Post-hoc analysis can extract dynamical motifs such as fixed points and line attractors, yet the ingredients that give rise to them remain implicit. More mechanistic models, constructed from first principles, offer greater interpretability, albeit at the cost of a narrower range of dynamics. For example, networks constructed from principles of global excitation-inhibition (EI) balance settle into stable background states, but their random connectivity cannot produce the complex spatio-temporal patterns of activity that accompany movement.^23–26^ Increasing the recurrent coupling of such networks can generate rich and sustained firing-rate fluctuations, but sacrifices stability and controllability.^27–30^

More recent mechanistic models exploit detailed EI balance in Dale-constrained connectivity to generate richer dynamics. A defining property of such Dalean connectivity matrices is that they are non-normal, i.e., their eigenvectors are non-orthogonal, enabling interactions between activity modes through hidden feed-forward cascades in which activity bleeds from one mode into another.^31–34^ A prominent feature of this non-normal structure is transient amplification, the temporary increase in network activity in response to a stimulus before it returns to baseline.^35^ Such models operate in a linear or quasi-linear regime: activity is amplified but not sustained, and producing persistent dynamics requires ongoing external drive.^31–33,36–39^

Here, we show that adding a single ingredient — neural non-linearity — to the requirements of stability and Dalean connectivity yields networks with a fully controllable repertoire of transient, steady-state, and periodic activity. We find that the interaction between transient amplification and neuronal non-linearity can push the system into a regime of self-sustained dynamics along a periodic orbit. To build such networks, we reverse-engineered the connectivity from a target Jacobian — the matrix that captures how neurons interact near the baseline firing rate. To construct the Jacobian, we selected a set of intrinsic dynamics — the eigenspectrum — and used the Schur decomposition to find the couplings between modes that render the connectivity Dalean. Our reverse-engineering approach yielded entire families of networks that require no hand-tuning of weights.

Without fitting them to neural data, our networks reproduced the experimentally reported population-level activity signatures of the motor cortex.^1,6,8,9,40,41^ The richness of cortical dynamics need not, therefore, rely on elaborate construction manuals or sophisticated plasticity rules; it may emerge from ubiquitous network properties such as Dalean connectivity, stability, and nonlinearity. Learning can then refine and orchestrate these dynamics rather than assemble them *ab initio*.

## Results

We approximated motor cortex dynamics using a recurrent neural network

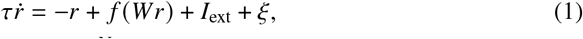

where *r* ∈ ℝ^*N*^ is the population activity with respect to baseline, *W* ∈ ℝ^*N* ×*N*^ is the connectivity matrix, and *f* (·) is the neuronal nonlinearity applied elementwise. *I*_ext_ ∈ ℝ^*N*^ represents external input and *ξ* ∈ ℝ^*N*^ is background noise. Unless otherwise stated, *I*_ext_ = 0, i.e., the network operated autonomously, reminiscent of a spring-loaded box that was triggered by a brief input and left to evolve freely.^32^ For simplicity, and without loss of generality, we set *τ* = 1 and chose *f* to be a shifted sigmoid such that *r*^∗^ = 0 is the baseline fixed point for all parameterizations (Methods).

We constructed networks that satisfy three conditions. The first is local stability: following any small excursion, the linearized dynamics near baseline must decay. This is determined by the Jacobian *J* = *f*′(0)*W* − *I*_*N*_, which approximates how neurons influence one another for small departures from rest. Local stability requires that all eigenvalues of *J* have negative real parts – in other words, every mode of activity must eventually decay. The second condition is Dale’s law: each neuron is either excitatory or inhibitory. Because Dalean connectivity is inherently non-normal, the third condition, transient amplification, is easily attained with sufficiently strong recurrent connectivity. Activity can then transiently grow in response to a stimulus before stability eventually returns the system to baseline.

To build a network that satisfies these conditions, we began by explicitly constructing Jacobians that are stable, Dalean, and amplifying, and derived their corresponding connectivity *W* = (*J*+*I*_*N*_)/*f*′(0). This reverse-engineering approach gives us direct control over the key ingredients – the amplification strength and how close a network sits to the edge of instability – allowing us to investigate how they interact with the neuronal nonlinearity to shape network dynamics. Notably, this approach requires no further training or hand-tuning of connectivity. For a small network with *N* = 2 neurons, the space of stable, Dalean Jacobians allows a closed-form characterization^31,38^ (Methods). For larger networks where no such characterization is possible (e.g., *N* = 100), we developed an algorithm that constructs Jacobians from a specified set of eigenvalues, and uses the Schur decomposition to find the couplings between eigenmodes such that the resulting connectivity is Dalean (Methods; see also Fig. 3A).

### Non-normality and nonlinearity ignite periodic orbits

With Dalean, stable, and amplifying networks at hand, we investigated whether persistent neural dynamics can emerge from the interaction between (i) non-normal transient amplification and (ii) nonlinear neural integration. In a reduced system with one excitatory and one inhibitory neuron, this interaction shaped the vector field into a stable periodic orbit that surrounded the baseline fixed point (Fig. 1B). The stable fixed point and the stable limit cycle coexist because the flow field contains a separatrix formed by an unstable periodic orbit (Fig. 1C, top). The geometry of this flow field is born from the directional structure imposed by the non-normality of *J* (Fig. 1C, bottom). Perturbations aligned with non-amplifying directions (*ϵ*_*T*_ ) decay back to baseline (Fig. 1D–E, bottom). By contrast, perturbations aligned with amplifying directions (*ϵ*_*O*_) experience transient linear growth near baseline. If the trajectory enters the nonlinear regime, nonlinear contributions then further sustain activity (Fig. 1D–E, top).

To better understand the interaction between amplification and nonlinearity, we decompose Eq. (1) into a linear and a nonlinear component by adding and subtracting the term *f*′(0)*Wr*:

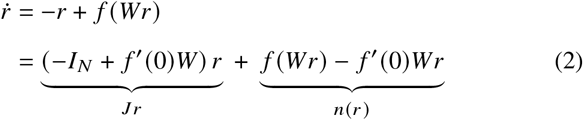

The linear term *Jr*, governed by the Jacobian, captures the dynamics near baseline. The nonlinear remainder *n* (*r*) vanishes near baseline, but grows as activity departs from the linear regime. This decomposition reveals the temporal signature of a network perturbation (Fig. 1F). For perturbations in amplifying directions, the linear term becomes positive at the onset of dynamics, which is the hallmark of transient amplification. The nonlinear term follows with a short delay; it remains positive throughout the trajectory, and becomes dominant after the linear term decays (Fig. 1F, top). As such, the nonlinear contribution carries the dynamics beyond the initial linear burst, igniting and sustaining periodic activity along the flow field of the system. Nonlinear saturation enforces an outer boundary beyond which trajectories cannot escape, even for large initial stimuli (Supplementary). For non-amplifying perturbations, both linear and nonlinear components rapidly decay and activity returns to baseline (Fig. 1F, bottom). Together, our results reveal a simple dynamical principle: transient amplification displaces the system into a nonlinear regime, where nonlinear components align with directions of growth and sustain persistent activity along a periodic orbit.

### A reservoir for complex persistent dynamics

We asked whether the interactions between linear and non-linear components could be the backbone for a repertoire of controllable, persistent activity patterns in larger cortical networks. We therefore extended our analysis to a network of 80 excitatory and 20 inhibitory neurons. Its Dalean connectivity (Fig. 2A,B) was constructed such that it was non-normal, had a stable baseline, and the excitatory and inhibitory synaptic weights onto each neuron summed to approximately zero (Fig. 2C; Methods). The resulting dynamics following a brief input (Fig. 2D) were reminiscent of cortical activity observed during movement.^1,6^ The direction and amplitude of these inputs sufficed to control duration and persistence of rhythmic and goal-directed movement dynamics, without additional sustained drive (Fig. 2D; Methods).

**Figure 2.**
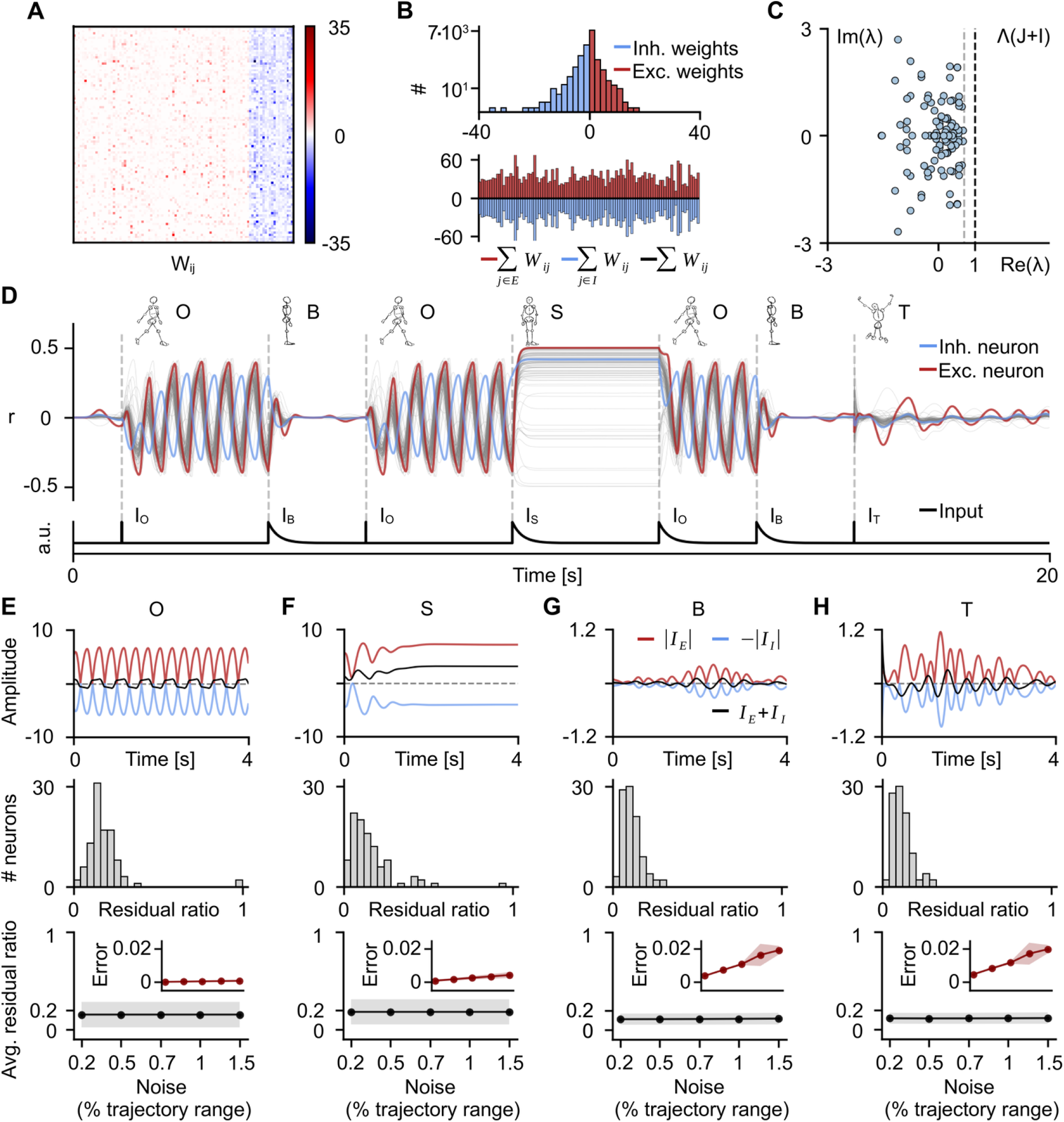
Brief transition commands control persistent dynamics in large-dimensional Dalean networks. **A)** Dalean connectivity matrix *W* with 80 excitatory (red) and 20 inhibitory (blue) neurons. **B) Top:** Distribution of all excitatory (red) and inhibitory (blue) synaptic weights of the network. **Bottom:** Cumulative excitatory (red) and inhibitory (blue) synaptic strength onto each neuron. The black line denotes the sum of all synaptic strengths. **C)** Eigenspectrum of the Jacobian *J* shifted by the identity matrix (Methods). Eigenvalues are confined to the stable regime, left of the (dashed) stability line and the imposed spectral abscissa (*α* = 0.7). **D)** Population dynamics are controlled by a series of brief transition commands. Amplifying directions (*I*_*O*_) initiate self-sustained oscillations (O). Non-amplifying directions (*I*_*T*_ ) generate transient responses (T). Additional inputs (*I*_*B*_ and *I*_*S*_ ) return activity to baseline (B), or drive the system toward non-trivial steady states (S). Red and blue traces show an example excitatory and inhibitory neuron; gray traces denote the remaining 98 neurons. **E–H) Top:** Excitatory (red) and inhibitory (blue) synaptic currents received by a neuron in the four dynamical regimes (B, T, O, S). The black trace shows the residual current (the instantaneous sum of excitatory and inhibitory inputs). **Middle:** Distribution of the time-averaged normalised residual current (residual ratio) across neurons. **Bottom:** Mean residual ratio as a function of background noise amplitude, averaged across neurons and noise realizations. **Inset:** Root mean square deviation between trajectories with and without noise.

For example, as in the reduced system, inputs that were aligned with amplifying directions (*I*_*O*_) drove the network away from baseline. The interaction between non-normal amplification and the saturating nonlinearity then triggered self-sustained oscillations (Fig. 2D,O). Inputs aligned with non-amplifying directions (*I*_*T*_ ) produced transient responses (Fig. 2D,T) that decayed back to baseline. In a large network, the higher dimensional vector field allowed for additional distinct, stable steady states (Fig. 2D,S), further enriching the dynamical repertoire. Oscillatory, transient and steady-state dynamics could thus coexist within a single network, controlled entirely by brief external commands (*I*_*O*_, *I*_*B*_, *T*_*S*_, *I*_*T*_ ). Activity could be initiated and stopped (Fig. 2D,B) *ad libitum*, and transitions between dynamical states occurred smoothly.

Across this repertoire of dynamics, the network operated in the loosely balanced excitation/inhibition regime characteristic of cortex.^42,43^ The instantaneous residual current — the sum of excitatory and inhibitory synaptic inputs onto a given neuron — remained small (Fig. 2E–H, top). The residual ratio — the time-averaged residual current relative to the average size of total recurrent input — indicated that the residual current was comparable to, though smaller than, the individual excitatory and inhibitory inputs (Fig. 2E–H, middle). This *loose* balance was maintained not only at baseline (B) but along the periodic orbit (O), at non-trivial steady states (S), and during transients (T), and was preserved across noise levels (Fig. 2E–H, bottom). Deviations between noisy and noiseless trajectories remained small across all four dynamical regimes, but not uniformly so: the baseline fixed point was the most susceptible to perturbation, i.e., departures from baseline and, as such, the initiation of movement-related activity, were easily attained. The periodic orbit and non-trivial steady states, being strongly attracting, exhibited the greatest robustness (Fig. 2E–H, bottom inset). The interaction between transient amplification and neuronal nonlinearity thus determined the existence or absence of these dynamical states, as well as how strongly the network was drawn to each one of them.

### Amplification and nonlinearity jointly shape the dynamical landscape

To understand better how the dynamical regimes described above are shaped by the interplay between the network phenomenon of amplification and the cellular property of nonlinearity, we systematically varied each ingredient while holding the other fixed. We first held the nonlinearity fixed and varied the homogeneity and local density (clustering) of the eigenvalue spectrum used to construct W. Although amplification is not strictly determined by the eigenspectrum, it varies with eigenvalue clustering in non-normal networks of the kind considered here (Fig. S1).^33,35^ Using our algorithm (Fig. 3A; Methods), we constructed three ensembles of 1000 networks whose Jacobians had eigenvalues sampled from distributions with low, medium, and high clustering (Fig. 3B; Methods). For each network, we computed the maximum transient amplification of the linear term *Jr*, defined as the largest peak in firing rates across 100 orthogonal initial conditions of fixed magnitude (Methods). We observed that amplification increased with clustering by several orders of magnitude, ranging from ∼10^1^ at low clustering to ∼10^8^ at high clustering (Fig. 3C, blue shades).

**Figure 3.**
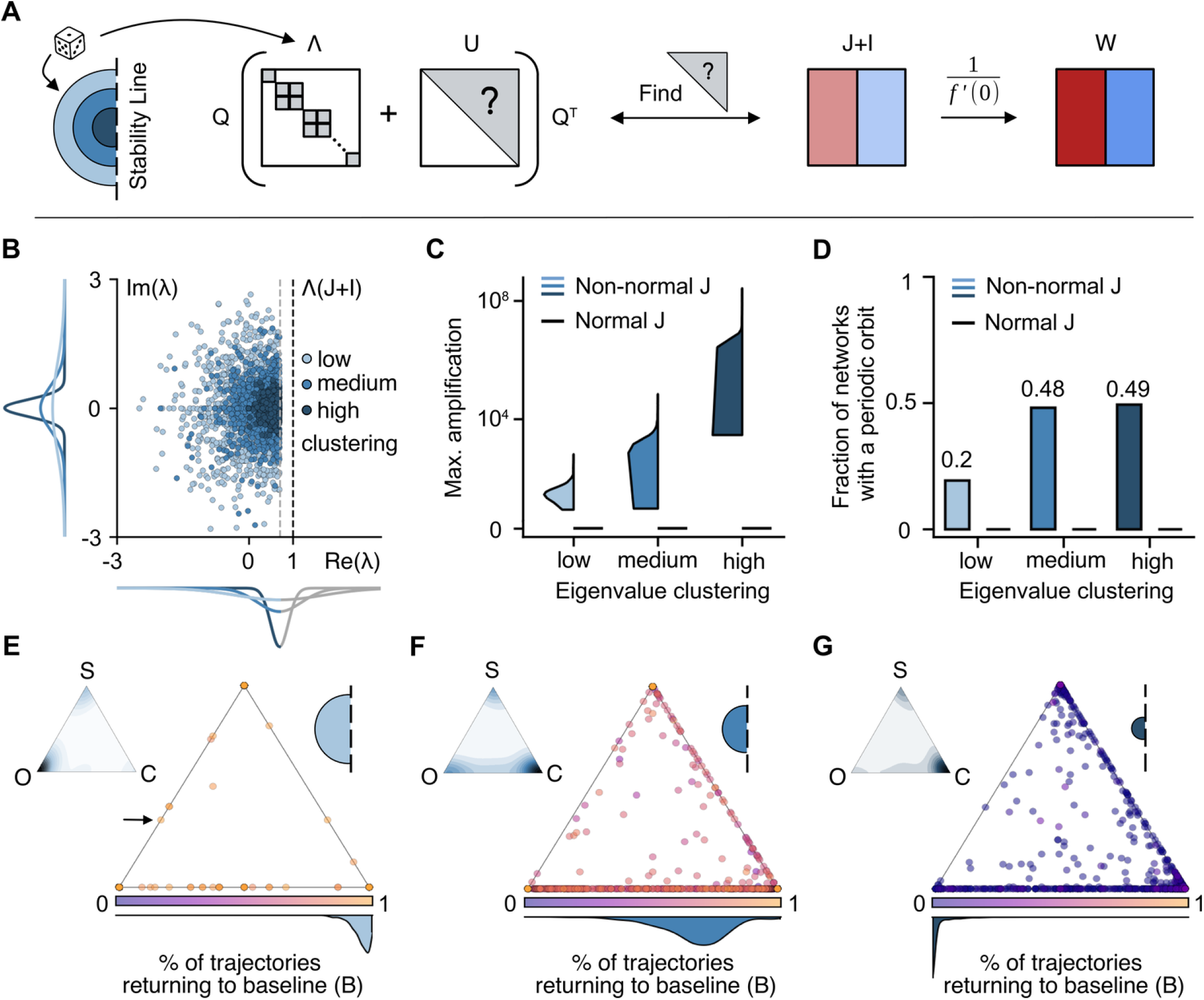
Higher amplification yields orbits, steady states, and chaos beyond baseline. **A)** Conceptual schematic of the connectivity matrix *W* construction. Stable eigenvalues are sampled from a desired distribution to form the normal (spectral) component Λ of the system. The normal component, together with an unknown non-normal component *U* and a unitary basis *Q*, forms the real Schur decomposition of the shifted Jacobian (*J*+*I*; Methods). The non-normal component *U* is determined by solving a set of linear inequality constraints such that the resulting *J*+*I* is Dalean. The connectivity matrix *W* is a rescaling of *J*+*I* that preserves Dale’s law. **B)** Complex eigenspectra of the *J*+*I* for three ensembles with low (light blue), medium (mid blue), and high (dark blue) eigenvalue clustering (1000 networks per ensemble; *N* = 100), as sampled from the three distributions plotted along x and y axis. Eigenvalues are confined to the stable regime, left of the (dashed) stability line and the imposed spectral abscissa of *α* = 0.7. **C)** Distribution of the maximum transient amplification (log scale; Methods) as a function of the clustering of eigenvalues for non-normal Jacobians (blue) and normal (control) Jacobians (black). **D)** Fraction of networks whose nonlinear dynamics exhibit at least one periodic orbit across 100 orthogonal initial conditions (Methods), shown for non-normal (blue) and normal (black) J. **E–G)** State occupancy across initial conditions for each network in the low (E), medium (F), and high (G) clustering ensembles. Each trajectory is classified as returning to the baseline fixed point (B), converging to a non-baseline steady state (S), approaching a periodic orbit (O), or exhibiting chaos (C). Each point represents a single network; normalised ternary coordinates show the relative fractions of S/O/C among non-baseline outcomes. Color indicates the fraction of trajectories returning to baseline (B). The distribution of baseline-return fractions across networks is plotted at the bottom. Inset triangles show a kernel-density estimate of the ternary distribution. The arrow in (E) indicates the example network used in Figs. 2, 6, and 7.

To discern how clustering — and hence amplification — shaped network behaviour, we classified the dynamics resulting from each of the initial conditions as returning to baseline (B), converging to a non-baseline steady state (S), approaching a periodic orbit (O), or exhibiting chaos (C). The fraction of networks with a periodic orbit rose sharply with clustering, from 20% in the low-clustering ensemble to approximately 50% in the medium-, and high-clustering ensembles. As expected, normal (control) Jacobians with identical eigenspectra showed no amplification at any clustering level (Fig. 3C, black) and never produced oscillations (Fig. 3D). State-occupancy plots (Fig. 3E–G) revealed how trajectory outcomes redistribute with clustering: at low clustering, most trajectories return to baseline, whereas at higher clustering, a growing fraction converge to steady states, periodic orbits, or chaotic attractors instead.

Next we held the connectivity fixed and varied the shape of the nonlinearity, running parameter sweeps for the gain *g*, threshold *θ*, and range *r*_*max*_ of the sigmoid function (Fig. 4A). The fraction of parameter combinations that could generate periodic orbits increased with amplification (Fig. 4B). Across eigenvalue clustering ensembles, these parameterizations concentrated in a band of low *θ* · *g*, largely independent of *r*_*max*_ (Fig. S2A-C). More strikingly, oscillations could be “rescued” in almost all networks across the three ensemble given the *right* nonlinearity (Fig. 4C, Discussion).

**Figure 4.**
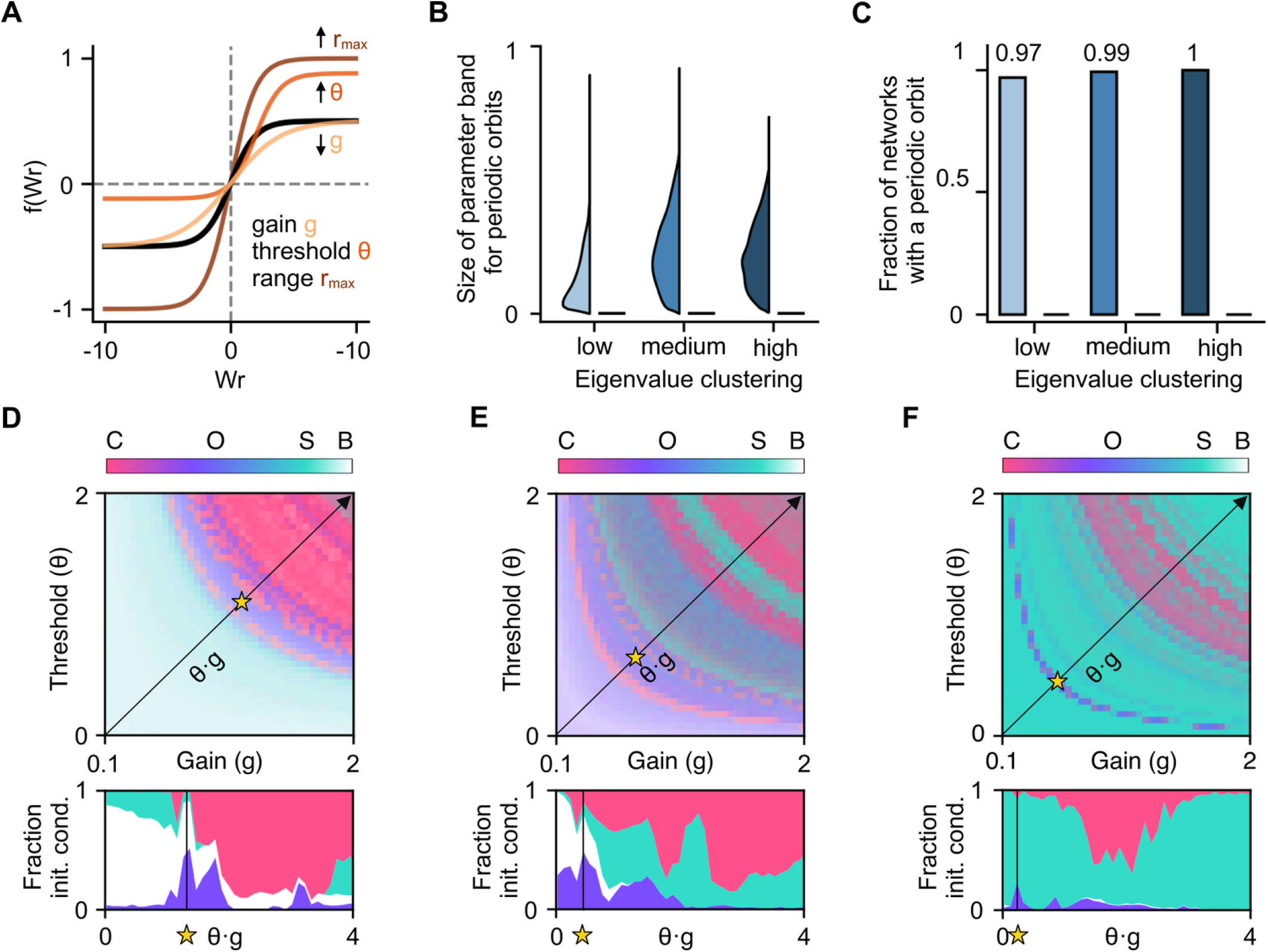
The shape of the nonlinearity redraws the dynamical landscape and can rescue periodic orbits. **A)** Schematic of the input-output function *f* (*Wr*) with the three parameters that shape it: gain *g* (gold), threshold *θ* (orange), and range *r*_*max*_ (brown), each varied around the default parameterization (black, *g* = 1, *θ* = 0, *r*_*max*_ = 1). **B)** Distribution across networks of the fraction of nonlinearity parameter combinations (gain *g*, threshold *θ*, and range *r*_*max*_ ) for which at least one initial condition converges to a periodic orbit, shown as a function of eigenvalue clustering for non-normal (blue) and normal control (black) Jacobians. **C)** Fraction of networks with at least one periodic orbit across nonlinearity parameter combinations, shown for non-normal (blue) and normal (black) J. **D–F)** Dynamics of one example network from the low (D), medium (E), and high (F) clustering ensembles as a function of nonlinearity parameters. **Top:** for every combination of gain (*g*) and threshold (*θ*), each network is initialized with 100 orthogonal initial conditions (Methods). Color indicates the fraction of trajectories converging to chaos (C, pink), a non-baseline steady state (S, teal), or a periodic orbit (O, purple); saturation indicates the fraction returning to baseline (B). **Bottom:** fraction of initial conditions converging to each dynamical state (B, O, S, C) as a function of the product *θ* · *g*.

To dissect how the nonlinearity determines the fate of a trajectory, we examined how the distribution of attractors changed with gain *g* and threshold *θ* for one example network from each clustering ensemble (Fig. 4D–F; increasing *r*_*max*_ only made the system slightly more linear, Fig. S2D–F). For the weakly amplifying network (Fig. 4D, top), most parameterizations supported a mixture of baseline returns, orbits, and steady states (light blue), with a band of predominantly oscillatory dynamics at intermediate nonlinearity strengths (purple) that gave way to chaos at the strongest parameterizations (pink). In the medium amplification example (Fig. 4E, top), the oscillatory band shifted toward weaker nonlinearities, baseline returns became less frequent, and the fraction of trajectories converging to steady states grew. In the strongly amplifying example (Fig. 4F, top), the network saturated rapidly across nearly the entire parameter range, converging predominantly to non-baseline steady states, with only a narrow band of orbits at the weakest nonlinearities, and almost no baseline returns. As dictated by our method of constructing connectivity matrices, dynamics depended on *g* and *θ* only through their product *θ* · *g*, so trajectory outcomes were symmetric along curves of constant *θ* ·*g* (Methods). Collapsing the parameter grid along this diagonal yielded a one-dimensional summary in which the same progression of regimes was visible at a glance (Fig. 4D–F, bottom).

These results demonstrate that a single network can support multiple coexisting attractors, with amplification gating the escape from baseline and nonlinearity selecting which attractor a trajectory ultimately reaches. The distribution of states that a network can host shifts with the amount of amplification the non-normal structure provides, and with the strength of the nonlinearity. Excess amplification, which would otherwise drive trajectories into saturation or chaos, can be offset by weakening the nonlinearity. Limited amplification can be compensated for by strengthening the nonlinearity. Stronger amplification at fixed nonlinearity, and stronger nonlinearity at fixed amplification, both drive the network through a similar sequence of regimes, where the two ingredients play partly substitutable roles, and jointly shape where the network sits within its dynamical landscape.

### Richer dynamics come at the cost of fragility

We wondered how forgiving the networks and their attractors were to perturbations in their connectivity matrices. To this end, we computed the perturbation margin *m* (*J*), the smallest change in connectivity that would catastrophically destabilise the network (Methods).^35^ We observed a trade-off between the perturbation margin and the ability of the network to transiently amplify (Fig. 5). Weakly amplifying, weakly clustered networks tolerated perturbations six to ten orders of magnitude larger than strongly amplifying, densely clustered ones (Fig. 5A). The origin of this trade-off became apparent in the *ε*-pseudospectrum, which maps the regions of the complex plane where eigenvalues can be pushed by perturbations of a given size *ε* (Methods).^33,35^ For a weakly clustered example, only the largest pseudospectral contours crossed the stability line (Fig. 5B, triangle). For a densely clustered example, even small contours extended deep into the unstable half-plane (Fig. 5C, square). Both example networks could ultimately be destabilised, but they differed by orders of magnitude in the perturbation size they could absorb before catastrophe happened.

**Figure 5.**
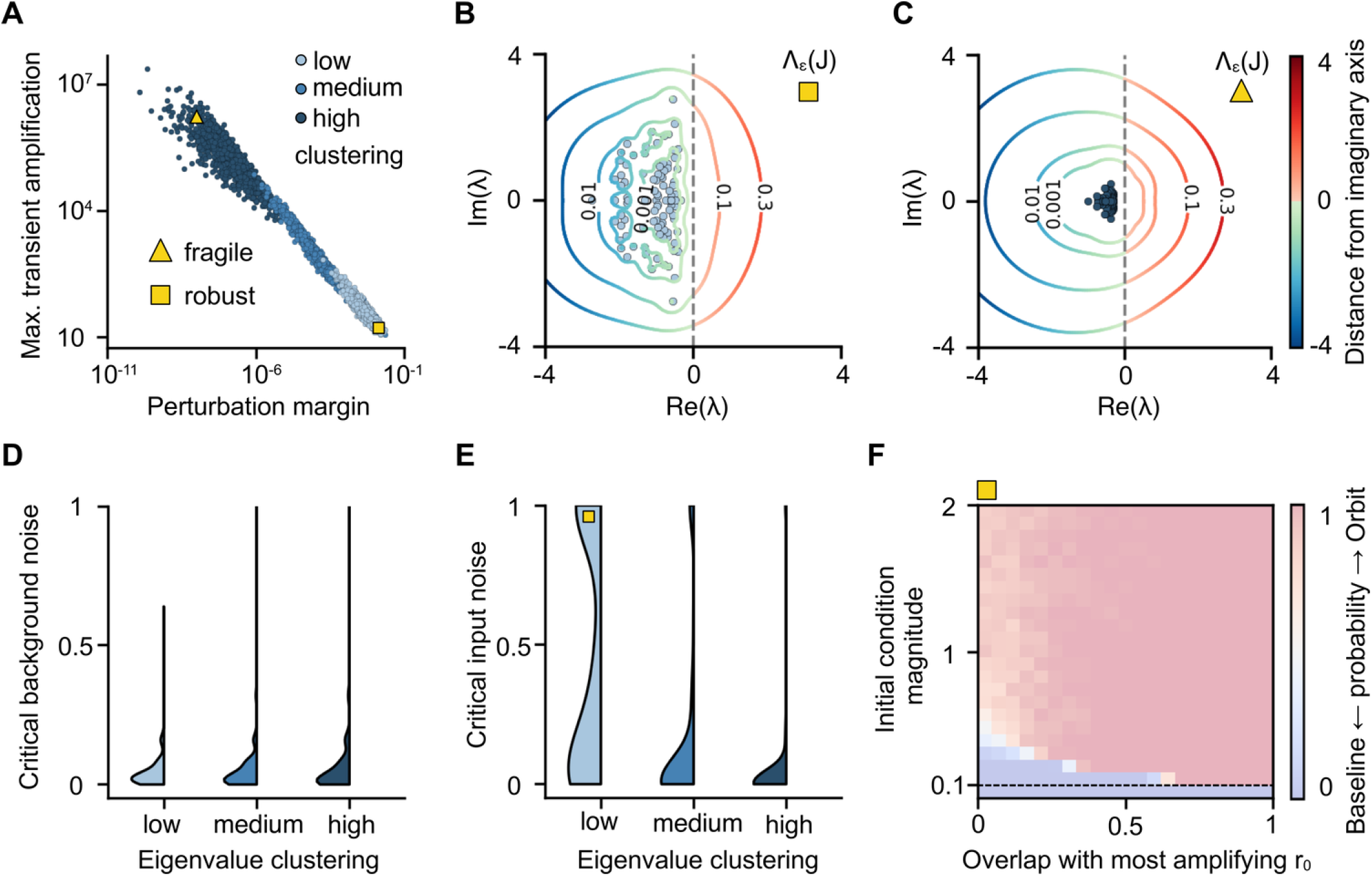
Trade-off between amplification and robustness. **A)** Maximum transient amplification plotted as a function of the perturbation margin *m* (*J*) of the connectivity matrix (Methods), for low (light ), medium (mid ), and high (dark blue) eigenvalue clustering of individual Jacobians (*J*). Yellow triangle and square denote two example Jacobians that produce robust and fragile dynamics, respectively. Axes are shown in log scales. **B-C)** Pseudospectra of the robust (B) and fragile (C) examples in (A). Contours indicate the maximum expected displacement of eigenvalues (circles) for perturbations of increasing operator norm (*ε*). Shades of red indicate where perturbations push eigenvalues into unstable regions, right of the (dashed) stability line, whereas shades of blue indicate stable regions. The colorbar shows the signed distance from the stability line. **D-E)** Distributions across networks of critical background (D) and input (E) noise beyond which trajectories diverged from the periodic orbit (Methods). **F)** Probability of converging to the periodic orbit (pink) versus returning to baseline (blue) as a function of initial condition magnitude and overlap with the most amplifying direction *r*_0_. The dashed line indicates the approximate boundary of the linear regime used in the stability analysis.

We also wanted to understand how perturbations to the dynamics themselves affected the behaviour of the networks. We therefore applied varying amounts of background and input noise to every network supporting a periodic orbit. We determined the critical amplitude of background noise above which trajectories escaped a stable orbit, as well as the critical amplitude of input noise beyond which trajectories failed to reach the orbit altogether (Methods). All networks, regardless of eigenvalue clustering, could tolerate comparable amounts of background noise (Fig. 5D), indicating that orbits acted as strong attractors. Conversely, input noise sensitivity varied considerably with clustering: networks with strongly clustered eigenvalues proved less tolerant to noisy initial conditions, while more weakly non-normal networks could tolerate a wider range of perturbations (Fig. 5E). The probability of converging to a stable orbit versus returning to baseline was shaped jointly by the magnitude of the initial condition and its overlap with the most amplifying direction *r*_0_ (Fig. 5F). Together, these findings suggest that the non-normal structure underlying periodic orbits must be close enough to instability to amplify, yet far enough to sustain robust dynamics.

### Gain modulation as a dial for oscillation frequency

Dynamics should be robust, yet not rigid in order to respond to changes in the environment.^44–46^ A useful motor network must thus adapt the properties of its periodic orbits when necessary. A natural candidate for such control is neuromodulation, thought to shape circuit computation by altering the input–output sensitivity of individual neurons.^47–50^ Similar mechanisms have been postulated to support flexible dynamics in the motor cortex.^17^ So we wondered whether neuromodulation could mold the frequency and shape of a periodic orbit.

We applied gain modulation to excitatory neurons, i.e., we changed the slope *α* of their activation functions (Fig. 6A; Methods). Such changes in *α* allowed for a range of oscillation frequencies in the dynamics between 0.8 and 2 Hz, as also observed experimentally (Fig. 6B).^8^ Single-neuron waveforms changed smoothly with *α* for excitatory, as well as for the not directly modulated inhibitory neurons (Fig. 6C). Unlike rescaling of the membrane time constant, which would dilate time without altering shape (not shown), gain modulation changed both the duration and the waveform of single-neuron responses, consistent with experimental findings.^8^ To visualize the resulting family of orbits geometrically, we projected the population activity onto its first two principal components and identified a speed axis along which oscillation frequency varied monotonically (Fig. 6D).^8^ The orbits were stacked along this axis, which faster oscillations traversed at higher speed, as reported experimentally^51^ and in previous rate-network models.^8,15,18^ Beyond stacking, slower orbits also tilted away from the plane of the faster orbits, gradually occupying new dimensions of the activity space. Such tilting has been proposed as a way for the network to reshape motor output without abrupt transitions between solutions.^8,52^

**Figure 6.**
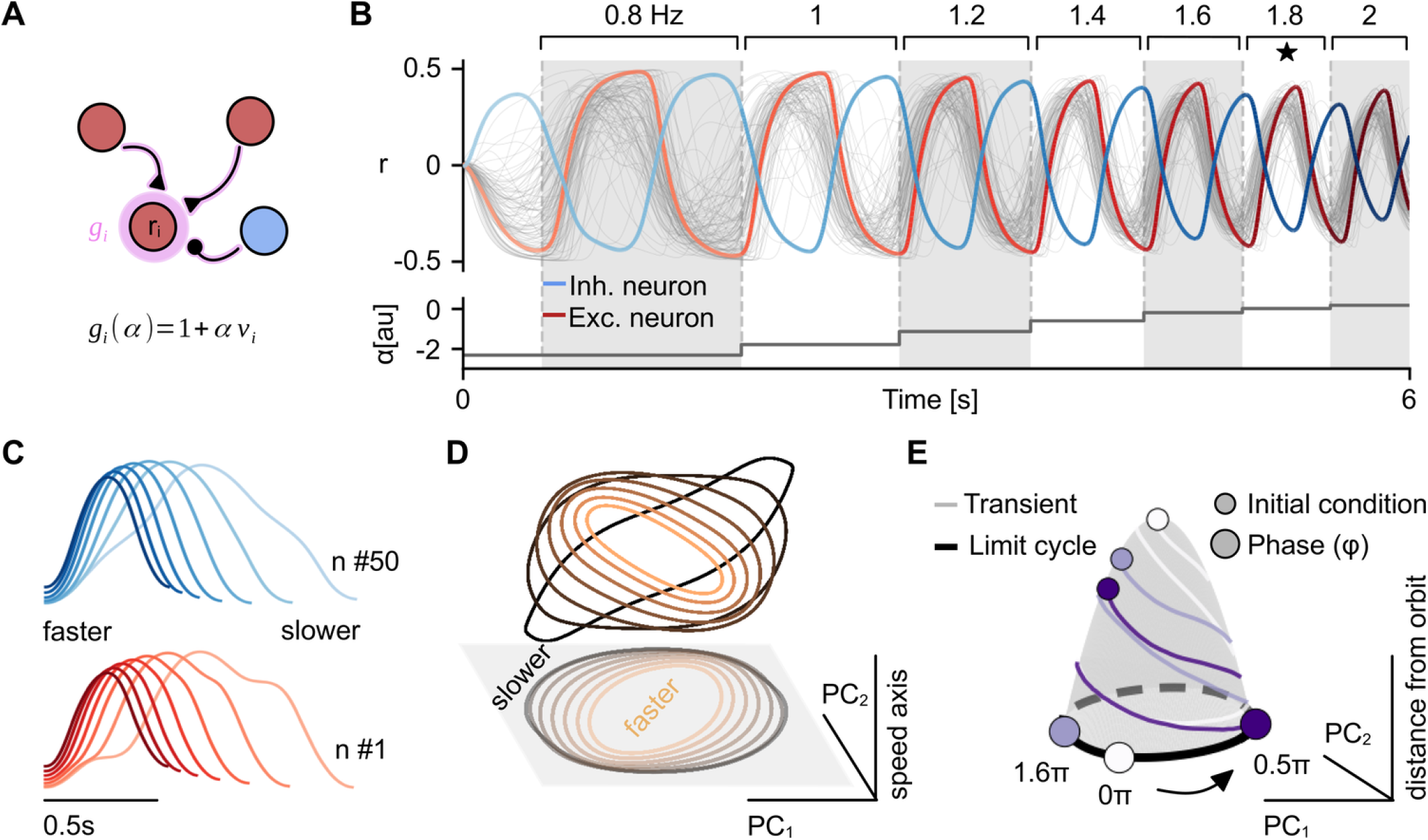
Oscillation frequency is smoothly controlled with gain modulation. **A)** Illustration of gain modulation. The input–output function of each excitatory neuron is scaled by a factor *g*_*i*(_*α*), modulating the strength of the total synaptic input it receives, according to a scalar control parameter *α* (Methods). **B)** Stepwise changes in the gain-control parameter *α*(*t*) (bottom, black) smoothly modulate the network’s oscillation frequency. Colored traces show example inhibitory (blue) and excitatory (red) neurons; gray traces denote the remaining 98 neurons. Frequency labels above each segment indicate the empirically measured oscillation frequency, spanning 0.8–2 Hz. The star marks the baseline gain condition *g* = **1** (i.e., *α* = 0). The network is the same as in Fig. 2. **C)** Activity waveforms of an example inhibitory (top, blue) and excitatory (bottom, red) neuron across values of the control parameter *α*, ranging from higher-frequency/faster (dark) to lower-frequency/slower (light). **D)** Projection of trajectories onto the first two principal components (PC_1_, PC_2_) together with a speed axis computed as in Saxena et al. (2022). Trajectory color indicates speed (orange: faster; brown: slower). **E)** Convergence to the periodic orbit from different initial conditions indicated by colour. The orbit is shown in a three-dimensional embedding (PC_1_, PC_2_, and distance-from-orbit). Small circles denote different initial conditions, which generate transients (gray surface) of varying duration before converging to the same orbit (black). Large circles indicate the phase *φ* at which trajectories approach the orbit.

We also observed that trajectories followed different paths before converging to the same periodic orbit at phases that depended on the starting position of the dynamics (Fig. 6E). Periodic orbits in our networks were thus malleable along multiple axes — frequency, waveform, and entry phase, mirroring the flexibility observed in motor cortex during rhythmic movements.

### Flexible motor sequences and context dependent dynamics

Motor cortex flexibility is not limited to shaping a single periodic orbit. The same network can produce qualitatively distinct rhythmic and goal-directed patterns of activity.^1,6,12,17,39^ Forward and backward cycling of the same limb, for instance, are represented by well-separated population trajectories within the same circuit.^6^ Our networks have so far produced only one periodic orbit alongside non-baseline steady states and multiple transients. We therefore asked whether a single network could host multiple periodic orbits and switch between them, without changing the connectivity. We show that contextual inputs, modeled as a constant per-neuron bias current *β* (Fig. 7A), can reshape the vector field and carve out new attractors without changing the connectivity of the network. Different biases could thus act as contextual signals that select different dynamical states (Fig. 7B). For example, in the 100-neuron network used in Figs. 2 and 6, five different bias currents *β* produced five distinct periodic orbits (Fig. 7C,D). These orbits occupied well-separated regions in activity space. Each context recruited activity in qualitatively different ways, with neuronal activity patterns differing across orbits in amplitude, phase, and shape (Fig. 7C). Pairwise distances between orbits, normalised by intrinsic orbit size (Methods), confirmed a clean separation rather than mild deformations of a common template (Fig. 7D).

**Figure 7.**
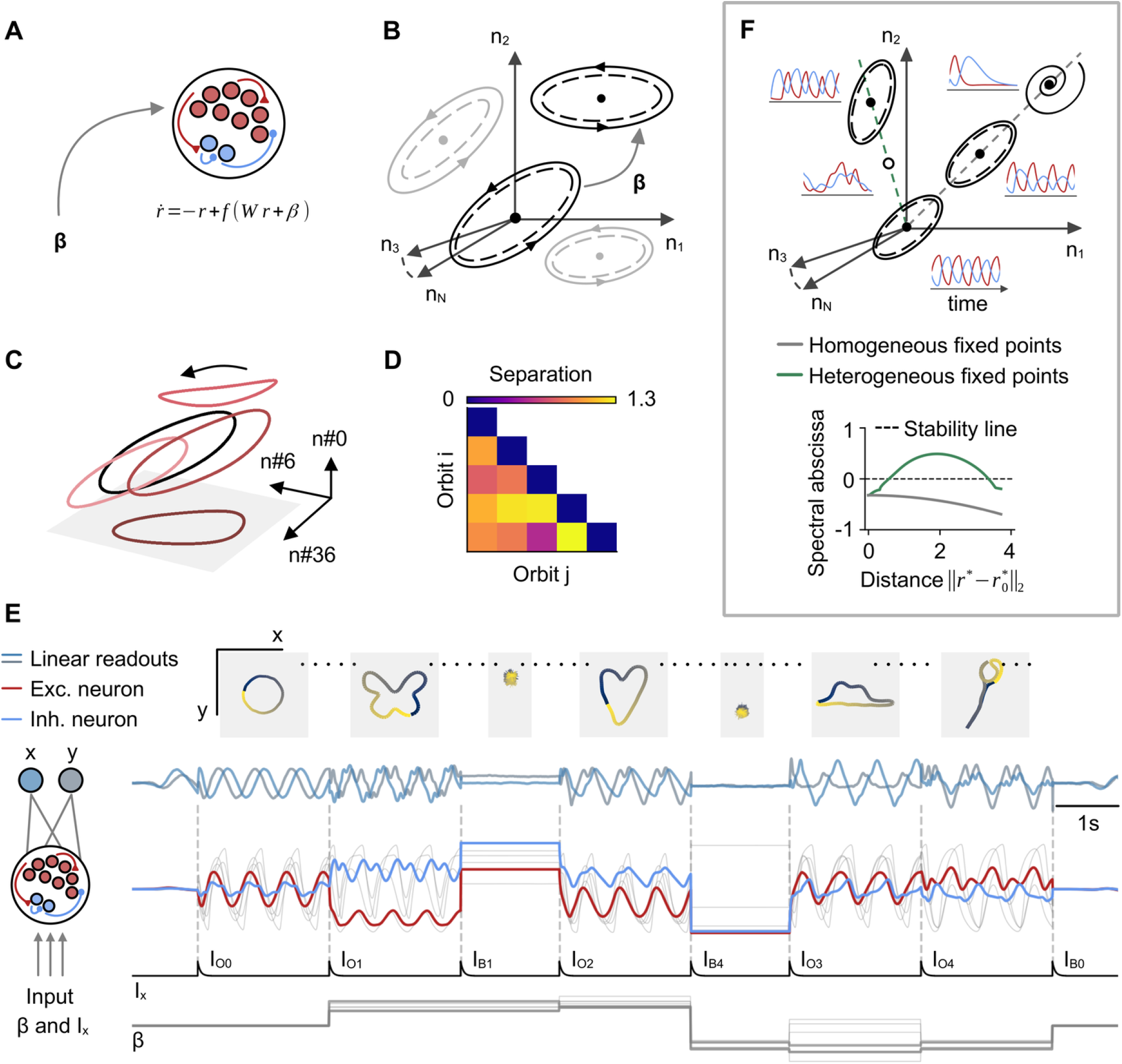
Contextual inputs carve out a repertoire of dynamics, including multiple orbits, within one network. **A)** Context is modeled as a constant per-neuron input, *β*. **B)** State-space illustration of how changing context *β* shifts the baseline fixed point (center) to a new position, and thus yields a new stable periodic orbit (solid), separated by an unstable orbit (dashed; Methods). **C)** Trajectories of 3 example neurons across 5 example context-dependent periodic orbits for the same network as in Figs. 2 and 6. **D)** Pairwise orbit separation, normalized by orbit radius; brighter colours indicate higher separation (Methods). **E)** Context-dependent neural activity patterns can orchestrate complex movement sequences. From bottom to top: contextual input (gray) and brief transition commands (black); 100 dimensional population activity (gray) over time, with example excitatory (red) and inhibitory (blue) neurons; two-dimensional (x, y) output decoded with a linear readout; the resulting trajectory mapped to x–y coordinates, with one traced shape per orbit cycle (Methods). **F)** Schematic of homogeneous and heterogeneous branches of bias-induced fixed points and their surrounding dynamics. **Top**: gray and green (dashed) lines indicate homogeneous and heterogeneous fixed-point manifolds, respectively; insets show the dynamics around stable (filled) and unstable (open) fixed points. **Bottom**: spectral abscissa along the homogeneous (gray) and heterogeneous (green) manifolds as a function of distance from the origin.

To demonstrate how a context-dependent repertoire of dynamics could provide the building blocks for motor sequences, we constructed a time varying contextual input *β*(*t*) that selected the attractor space, while brief transition commands *I*_*x*_ controlled the type and onset of dynamics (Fig. 7E, bottom; Methods). Together, *β* and *I*_*x*_ drove the network through a sequence of periodic orbits (Fig. 7E, red and blue activity traces); transients could be interleaved just as easily, and the number of embedded orbits could be expanded (not shown). A fixed two-dimensional linear readout trained jointly across all states^17,32^ (Methods) mapped population activity to motor output, producing one fully-traced shape per orbit cycle (Fig. 7E, top). A single connectivity matrix could thus generate a diverse and flexible ensemble of motor outputs, controlled entirely by slow contextual and intermittent, brief transition inputs.

To reveal how context-dependent inputs could enrich the dynamical repertoire of the network so dramatically, we investigated how a bias current *β* could create a new fixed point away from the origin. We found that the stability of the newly emerging fixed points, together with the geometry of the surrounding vector field, depends critically on how the bias is distributed across neurons (Fig. 7F; Methods). If all neurons receive identical *β*, the new fixed points align along a continuous, stable branch, with increasingly contracting dynamics further from the origin (Fig. 7F, top, gray). The spectral abscissa of the linearization decreases monotonically along this branch, so the local amplification that sustains periodic activity fades with distance, and homogeneous biases cannot produce new orbits far from baseline (Fig. 7G, bottom, gray). Heterogeneous biases on the other hand, where each neuron receives its own input, behave very differently. Stable fixed points appear as disconnected “islands” surrounded by unstable equilibria and chaotic dynamics (Fig. 7G, top, green). Any continuous path from baseline to such an island must cross unstable territory (Fig. 7G, bottom, green). Within an island, however, the local linearization remains amplifying and can host a new periodic orbit. These results show that contextual inputs can vastly increase the repertoire of network dynamics, both transient and persistent.

### Low dimensional trajectories tile a high dimensional repertoire space

Given the diverse repertoire of dynamics we found, we wondered about the structure and dimensionality of the space it occupies. Experimentally, it has been observed that motor cortical activity unfolds along low-dimensional manifolds,^16,40,41,53,54^ and recent findings suggest that different tasks may recruit distinct, low-dimensional subspaces that together tile a higher-dimensional space.^9,55^

To place our model within this framework, we assembled a repertoire of 100 trajectories by sampling 50 contexts, each defined by a bias current *β*, and selecting from each context its periodic orbit and one arbitrary transient (Methods). Individual trajectories in the repertoire were well described by at most four principal components, while all orbits and all transients required 28 and 38 components, respectively, to explain 90% of the variance. The full repertoire of all trajectories (orbits and transients) required 42 components, an order of magnitude above the dimensionality of any single trajectory. Similarly, a within-context subset of one orbit and 49 transients needed 31 components (Fig. 8A). Individual trajectories, whether orbits or transients, thus explored only a small fraction of the dimensions spanned by the full repertoire.

**Figure 8.**
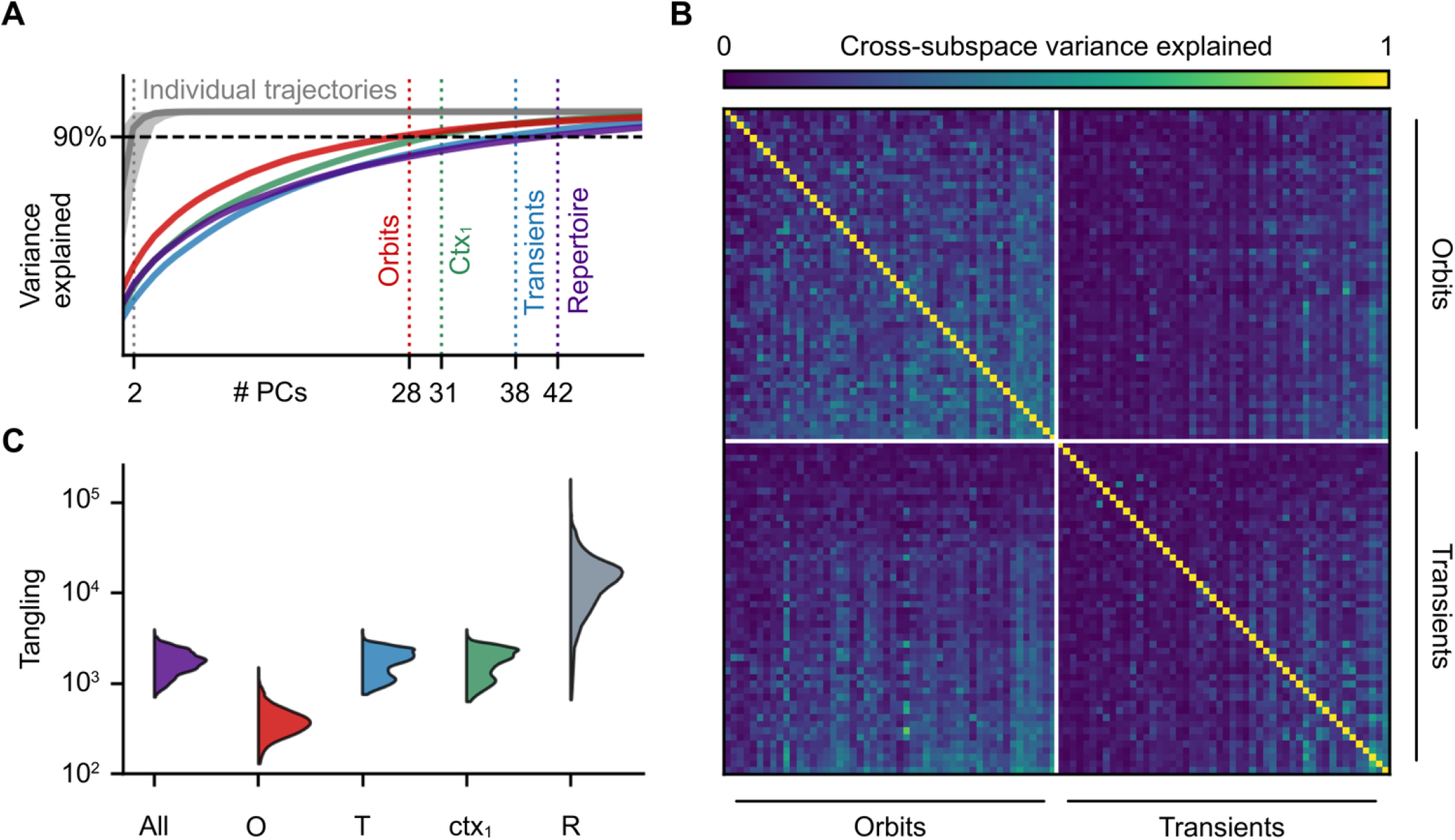
A high-dimensional repertoire of untangled, well-separated trajectories. **A)** Cumulative variance explained as a function of the number of principal components, computed for individual trajectories (gray line and shaded area show the mean and distribution across trajectories), the 50 orbits alone (red), the 50 transients alone (blue), a within-context set of one orbit and 49 transients (green), and the full repertoire (indigo, 50 orbits and 50 transients sampled across contexts). Dashed vertical lines mark the number of components that explain 90% of the variance for each subset. **B)** Pairwise cross-subspace variance among the 50 orbits and 50 transients of the repertoire. Each entry shows the variance of a target trajectory captured by the top-3 PCs of a given trajectory. Rows and columns are sorted by trajectory type (orbits, then transients). Colorbar (top) ranges from 0 (dark blue) to 1 (yellow). **C)** Distribution of tangling values (log scale), computed as in Russo et al. (2018) for the repertoire (All, indigo), orbits only (O, red), transients only (T, blue), trajectories within a single context (ctx_1_, green), and the two-dimensional linear readout (R, gray).

To understand how low-dimensional trajectories are distributed across the high-dimensional space, we computed the matrix of pairwise cross-subspace variance, which measures how well the leading principal components of one trajectory reconstruct another (Methods).^4^ Off-diagonal values of the matrix were close to zero across most pairs of orbits and transients (Fig. 8B). On average, the top three components of any one trajectory captured only 15% of the variance of another. This value was 20% within the orbit block, and 10% within the transient block. Trajectories therefore tile the repertoire space, occupying largely separate subspaces rather than crowding into a common one.

To further characterize the structure of these trajectories, we computed their tangling (Fig. 8C).^6^ A flow field is tangled when states close in neural space evolve along divergent directions, and untangled when nearby states move along similar paths. Tangling remained low across the full repertoire of 100 trajectories (Fig. 8C, indigo), within the orbit and transient subsets considered separately (Fig. 8C, red and blue), and within trajectories belonging to a single context, that is, sharing the same bias current *β* (Fig. 8C, green). These results are consistent with a smooth and robust flow field of the network (Eq. 1), where dynamics are shaped by internal recurrence rather than by external commands.^6^ The two-dimensional linear readout, by contrast, exhibited tangling at least an order of magnitude larger (Fig. 8C, gray), mirroring the experimental observation that muscle-like outputs are necessarily more tangled than the neural population activity that generates them.^6^

Overall, we show that a single recurrent network, driven only by brief transition commands and contextual inputs, can host a high-dimensional repertoire of low-dimensional, well-separated, untangled trajectories. This organization mirrors the population structure reported in motor cortex,^9^ and offers a mechanistic substrate for flexible motor sequences without training or hand-crafting of connectivity.

## Discussion

We have shown that three simple ingredients – Dalean connectivity, stability, and neural nonlinearity – suffice to reverse-engineer entire families of networks that produce a rich repertoire of transient, steady-state, and periodic activity, with no prior learning. The complex neural dynamics emergent in our networks come intriguingly close to those observed in primate motor cortex. Individual orbits and transients unfold, untangled, along low-dimensional manifolds,^6,16,40,41,51,53,54^ while the full repertoire collectively spans a higher-dimensional space, with different trajectories occupying largely non-overlapping subspaces.^9^ Our networks, akin to the motor cortex, thus behave like spring-loaded boxes, with population dynamics driven by internal recurrence rather than external drive.^1,2,6,32^ That these features emerge jointly, without fitting to neural data, suggests they share a common origin in the non-normal, nonlinear geometry of stable, Dale-constrained recurrence, which could also be the mechanism at work in the motor cortex.

### A distinct dynamical principle

The three ingredients that catalyze the emergence of mutlistable dynamics are well-grounded in cortical biology. Firstly, a stable activity baseline follows from the loose excitation/inhibition balance characteristic of the inhibition-stabilised regime in neuronal networks,^43,56–59^ and can be maintained by homeostatic regulation of firing rates.^60–62^ The second ingredient, non-normality, is generic for any Dale-constrained connectivity matrix,^31,33,42^ and the third, a saturating single-neuron response, arises from biophysical thresholds and ceilings on firing rates.^63^ How these ingredients combine to produce stable periodic orbits can be partly understood through analytics. We use the energy of the system to identify amplifying directions of outward drift near baseline and the conditions under which such drift appears. We show that firing-rate saturation imposes an outer boundary beyond which trajectories cannot escape (see Supplementary). Given the *right* choice of nonlinearity, all constructed networks could host a periodic orbit, emphasizing the ubiquity of the mechanism. The *right* choice of nonlinearity for each network may depend on how the linear and nonlinear components of the dynamics align at the onset of activity, as well as far away from baseline. The structural properties of the connectivity, i.e., how tightly balanced any given mode of activity may be, as well as how closely they overlap with other modes, set the magnitude and directions of amplification, and thus whether and when a trajectory can escape from baseline. The shape of the nonlinearity, in turn, selects which attractor the trajectory reaches farther from baseline. The interaction between the two determines the fate of the dynamics.

### Robustness

The simplicity of the interaction between non-normality and nonlinearity, together with the ubiquity of these ingredients, means that periodic orbits and, more broadly, multistable dynamics occupy a broad band of parameter space rather than a narrow ridge (Figs. 3,4,S3). Amplification and nonlinearity are partly interchangeable: weakening one can be compensated for by strengthening the other. The mechanism is thus robust and requires only coarse, rather than precise, control of either. Robustness of the mechanism, however, is distinct from robustness of the connectivity itself. The very features that produce the richest dynamics also push the network closer to the edge of instability, making particular instantiations of the connectivity more fragile to structural perturbations and noisy inputs, even though the orbits themselves act as strong attractors once reached (Fig. 5). The richest dynamics emerge from an intermediate band of solutions: too little amplification or nonlinearity, and trajectories return to baseline; too much, and they are driven into saturation or instability.

### Spiking networks

In this study we work within a rate-based framework, but the linear/nonlinear decomposition we use for rate dynamics has a direct interpretation in spiking networks. The linear term captures the propagation of small departures from baseline through the non-normal connectivity; in spiking terms, preferred input directions recruit more spiking activity from each neuron.^32,33^ The nonlinear remainder corresponds to supralinear firing at the single-neuron level, compounded over the population through recurrence. Co-tuned neurons reinforce each other through the non-normal couplings of the connectivity, and a periodic orbit emerges as a network-level phenomenon. No single neuron acts as an oscillator; the energy required to sustain the dynamics is supplied by spiking activity organised by network geometry. Rate-network predictions of this kind have been shown to transfer to spiking implementations,^64^ though a direct spiking realisation of the present construction remains to be realized.

### Experimental predictions

Our model produces distinctive signatures at two levels, both experimentally accessible: the geometry of the dynamical landscape, and the mechanism that shapes it. The coexistence of a stable fixed point and a stable periodic orbit in a single network implies a separatrix between the two basins of attraction. Adaptation notwithstanding, this geometry could be revealed in an experiment by varying the magnitude of the *right* (selective for the orbit) input. Small inputs should produce decaying responses, whereas inputs above a threshold should drive convergence to the orbit. Inputs near the threshold should produce long transients before settling on either attractor, reflecting the slow dynamics near the separatrix. These signatures pin down the geometry but not the underlying mechanism, which is the interaction between non-normality and nonlinearity. The Jacobian, which captures local linear interactions between neurons, could be estimated from sub-threshold population dynamics. The early phase of the response to an input that triggers self-sustained oscillations should be well captured by this linearization, and only later should the trajectory diverge from the linear prediction along directions shaped by the nonlinearity. Patterned holographic stimulation combined with simultaneous population recordings^65–69^ could provide the experimental handle for both geometric and mechanistic investigations.

### Feedback, learning and capacity

Our model operates in a mostly open-loop regime, without sensory feedback. At times, the network received slow, time-varying contextual inputs which reshaped the flow field and carved out new attractors without altering the connectivity (Fig. 7). Biologically, such inputs could originate from pre-frontal projections^14,70^, thalamocortical loops,^39^ or neuromodulatory systems.^50,71,72^ Faster, closed-loop control could likewise be incorporated through sensory feedback from the cerebellum and basal ganglia^73,74^ within the broader framework of optimal feedback control of cortical dynamics.^75,76^

Throughout the paper we have implicitly considered the network dynamics to be complete movements, such as arm reaches or cycling bouts. However, because context can in principle multiply the repertoire without bound, each trajectory could instead be interpreted as a motor primitive^17,77^ or subskill^9^ — a reusable element from which behaviour is composed. Under this view, the role of learning and plasticity shifts from sculpting dynamics out of an unstructured substrate, to shaping one that is rich to begin with. Synaptic changes in the motor cortex during learning^78,79^ could reshape existing primitives to meet changing task demands,^16,53^ or compositionally recombine them to arrive at regions of the activity space outside previously visited manifolds.^21^ Learning could also embed entirely new primitives, further expanding the repertoire.^37,55,80^ Together with mechanisms such as targeted gain modulation, which can silence functional subpopulations so that not every neuron participates in every movement,^17,20,81,82^ context-dependent dynamics and plasticity turn our networks into reservoirs with vast capacity, in the spirit of universal function approximators.^83^

## Conclusions

Computational models of motor cortex have typically embedded rich dynamics into networks by training them to reproduce observed activity. Our results invert this scheme. Three simple biological constraints — Dale-constrained connectivity, a stable baseline firing-rate, and neuronal nonlinearity — can form the backbone of emerging network dynamics. Under these constraints, networks can give rise to a rich repertoire of fully controllable transient and self-sustained dynamics. Such dynamics align with the population-level signatures reported in motor cortex during both goal-directed^1^ and rhythmic movements,^6^ and unfold along untangled, low-dimensional manifolds^40,54^ that collectively tile a high-dimensional space,^9^ without being fit to data. Gain modulation and contextual inputs further enrich the repertoire, granting the network the flexibility required for motor sequencing. The implication is conceptual as much as computational: richness need not be instilled in cortical circuits, because it is already what circuits obeying Dale’s law, stability, and nonlinearity posses. Learning and plasticity are then free to act on a substrate that is already expressive, rather than build one from scratch.

## Acknowledgments

This work was supported by the ERC Consolidator Grant (SYNAPSEEK). We thank Henning Sprekeler, Guillaume Hennequin, Everton Agnes, Basile Confavreux, Alexander Rivkind, Georgia Christodoulou, Jake Stroud, Panagiotis Bozelos, Veronica Scozzi and James Ferguson for insightful comments and discussions.

## Methods

### Network model

We consider a recurrent firing-rate network of excitatory and inhibitory neurons Miller and Fumarola, 2012:

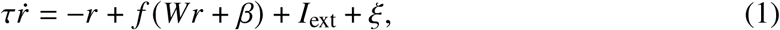

where *r* ∈ ℝ^*N*^ denotes the population activity with respect to baseline, *W* ∈ ℝ^*N*×*N*^ is the recurrent connectivity matrix, and *f*( ) is applied elementwise. The vector *β* ∈ ℝ^*N*^ represents a constant bias input, *I*_ext_ ∈ ℝ^*N*^ denotes brief control inputs, and *ξ* ∈ ℝ^*N*^ is an additive noise term. Unless otherwise stated, we set *β* = 0 and *I*_ext_ = 0.

All neurons share the same time constant *τ*. Because changing *τ* uniformly rescales time, the choice of *τ* does not affect the qualitative behavior of the system; we therefore set *τ* = 1 throughout without loss of generality.

The neuronal nonlinearity is a shifted sigmoid chosen such that *r*^∗^ = 0 is the baseline fixed point for all parameterizations:

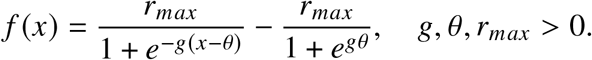

Close to the baseline fixed point, the dynamics of the deterministic system (*ξ* = 0) are well approximated by the linearization:

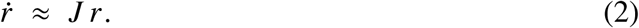

The fate of the linearized dynamics is determined by the eigenvalues of the Jacobian:

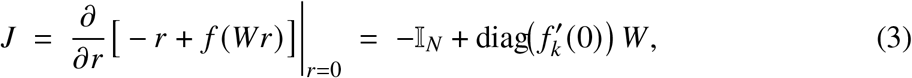

where *f*_*k*_ denotes the nonlinearity of neuron *k*. If all eigenvalues of *J* have strictly negative real parts, the fixed point is stable and activity returns to baseline following small perturbations.

Given a stable Jacobian, we then reverse-engineer the corresponding connectivity matrix:

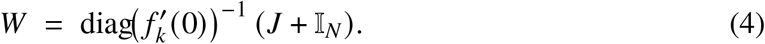

For a uniform nonlinearity across neurons, the connectivity matrix is *W* = (*J* + I_*N*_ )/ *f*′(0). With *r*_*max*_ = 1, the derivative of *f* at the fixed point simplifies as 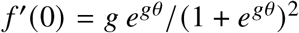, and the argument of the nonlinearity *f* (*Wr*) becomes:

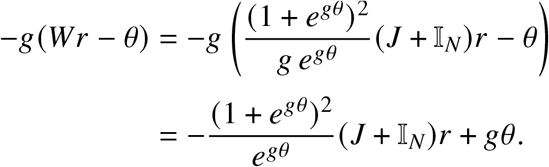

The nonlinearity is therefore parameterized by a single effective parameter *gθ*. The same cancellation holds neuron-wise for heterogeneous nonlinearities: when each unit *i* has its own pair (*g*_*i*_, *θ*_*i*_), the *i*-th component of the argument depends on these parameters only through the product *g*_*i*_*θ*_*i*_.

Note: dynamics can equivalently be expressed in strictly positive firing rates (Supplementary), and the baseline state need not be homogeneous across neurons (see Fig. 6).

### Transient amplification

Transient amplification occurs when the norm of the activity temporarily increases under linear(ised) dynamics (Bondanelli and Ostojic, 2020; Hennequin et al., 2014; Trefethen and Embree, 2005):

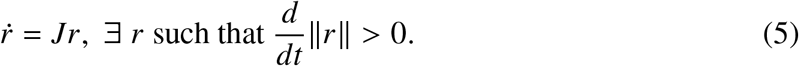

The maximum transient amplification of a system is defined as the largest increase in activity norm relative to the norm of the activity at time *t* = 0, evaluated across time and multiple initial conditions *r*_0_:

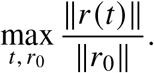

### Dalean, stable, amplifying Jacobians

We construct Jacobians *J* whose eigenvalues have negative real part and whose weights satisfy Dale’s law. Throughout, we work with the shifted matrix

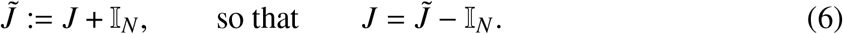

This parameterization is convenient because the underlying connectivity matrix *W* is recovered from 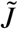 by a positive rescaling (at the fixed point), so Dalean sign constraints can be imposed directly on 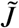 and are preserved under this mapping.

Stability of the fixed point requires ℜ(*λ*(*J*)) < 0, and equivalently

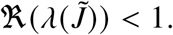

Dale’s law is enforced column-wise. Every presynaptic neuron *j* is either excitatory ( *j* ∈ *E*) or inhibitory ( *j* ∈ *I*), constraining the sign of its outgoing weights:

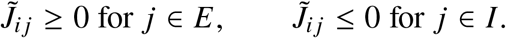

#### Two-dimensional Jacobians (closed-form filtering)

For *N* = 2, we adopt a construction similar to prior low-dimensional stability-amplification analyses (Bondanelli and Ostojic, 2020; Murphy and Miller, 2009). We write

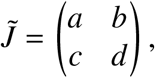

and impose Dale’s law with neuron 1 excitatory and neuron 2 inhibitory:

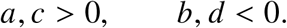

#### Stability

Let *λ*_1_, *λ*_2_ denote the eigenvalues of 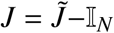. Local stability requires ℜ(*λ*_1_), ℜ(*λ*_2_) < 0. For a real 2 × 2 matrix this is equivalent to

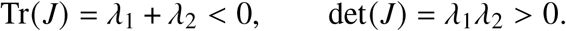

Using 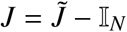,

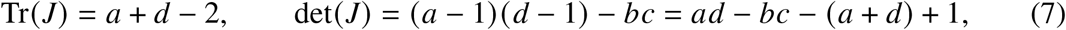

so stability is equivalent to

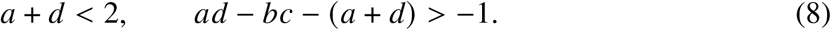

#### Transient amplification

Transient amplification is defined by 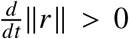 for some state *r* under the linearized dynamics 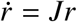. With 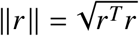,

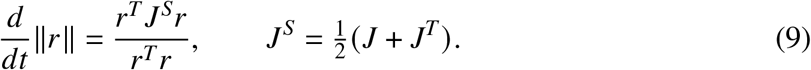

Amplification exists if and only if *λ*_max_(*J*^*S*^) > 0. Since *J*^*S*^ is symmetric and Tr(*J*^*S*^) = Tr(*J*) = *a* + *d* − 2 < 0 in the stable regime, this is equivalent to

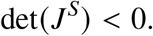

Substituting 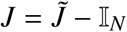 gives 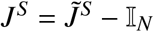 with

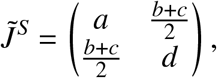

hence

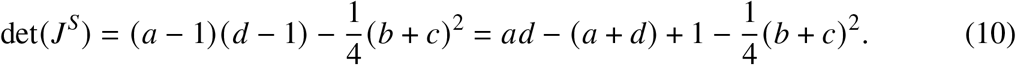

Using (*b* + *c*)^2^ = (*b* − *c*)^2^ + 4*bc*, the amplification condition can be written as

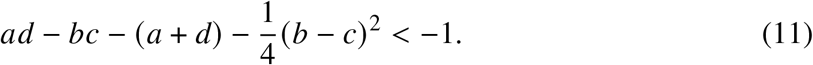

We sample (*a, b, c, d*) and retain only realizations satisfying Dale’s law and the stability and amplification inequalities, with an additional magnitude bound |*a*|, |*b*|, |*c*|, |*d* | ≤ 5.

#### Large dimensional Jacobians (Schur form and linear programming)

For *N* > 2, we specify the spectrum Λ of 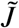 and solve for non-normal couplings *U* that enforce Dale’s law. We represent 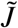 in real Schur form,

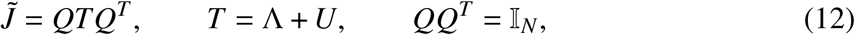

where Λ is block-diagonal (real Schur blocks) and *U* is strictly upper triangular, so that Λ fixes the eigenvalues of 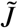 and *U* controls non-normality. 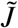 is thus split into a normal and a non-normal component:

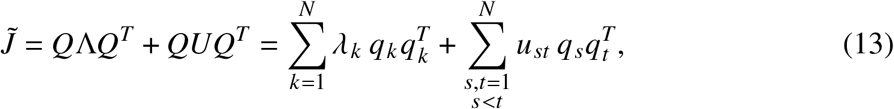

where *q*_*k*_ ∈ ℝ^*N*^ is the *k*-th column of *Q*, so that each 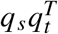 is a rank-one *N* × *N* matrix.

#### Sampling eigenvalues

We sample 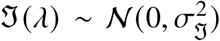 and ℜ (*λ*) from a normal distribution centered at a target spectral abscissa *α* and truncated to (−∞, *α*],

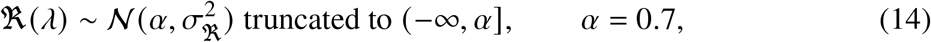

and enforce complex-conjugate pairing. Since 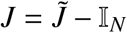, imposing 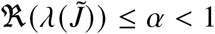 ensures ℜ(*λ*(*J*)) < 0.

#### Dale constraints as linear inequalities

For fixed *Q* and Λ, 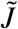 depends on the unknown upper-triangular entries of *U*; vectorizing these entries into *u* = vec(*U*), we write

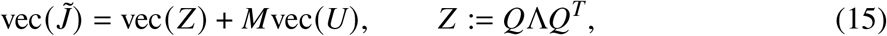

so Dale’s law becomes a system of linear inequalities in *u*, solved via linear programming.

#### Constructing *Q*

We initialize

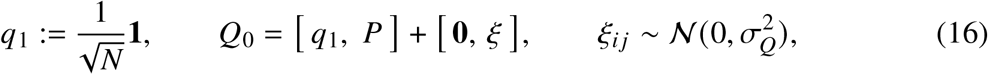

where *P* ∈ {0, 1}^*N*×(*N*−1)^ has a single value of 1 per column, in unique rows, and noise is added to all columns except the first. We then orthonormalize *Q*_0_ (Gram–Schmidt) to obtain an orthogonal matrix *Q* whose first column is *q*_1_.

Fixing 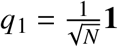 decomposes 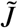 into a uniform baseline, a per-column offset, and *R*:

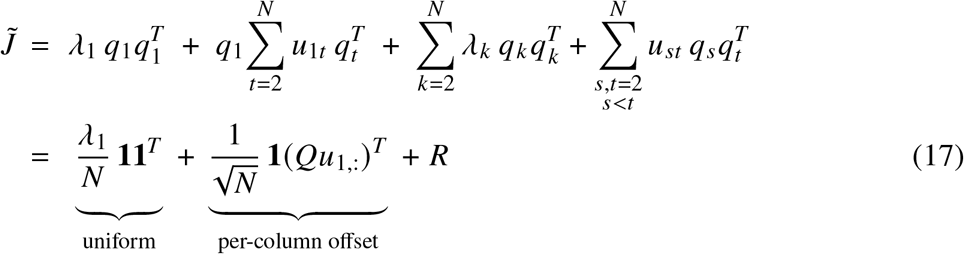

where *u*_1:_ = (*u*_11_, …, *u*_1*N*_ )^*T*^ is the first row of *U*.

The uniform component and the per-column offset are crucial in determining the feasibility of the system, i.e., whether, for fixed *Q* and Λ, we can find a Dalean 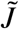. The uniform baseline adds *λ*_1_/*N* to every entry of 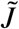, whereas the per-column offset shifts the baseline of each column independently—more positive for excitatory columns and more negative for inhibitory columns—thereby realising the Dalean constraint. With *e* _*j*_ the *j* -th standard basis vector and *m* = *Qu*_1:_,

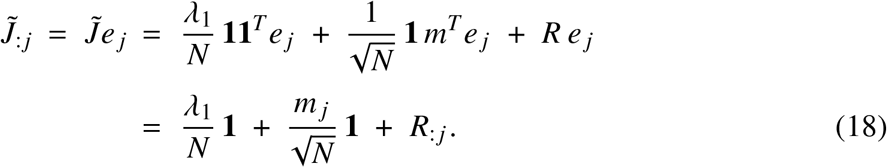

Because the columns *q*_2_, …, *q*_*N*_ are orthogonal to *q*_1_, they must have zero mean, and hence so does every column *R*_:*j*_ . *R*_:*j*_ therefore cannot shift the baseline of column *j* — which is already fixed by the uniform and per-column offsets — but, through the remaining free entries of *U*, it supplies the within-column corrections needed to satisfy Daleanity.

Here, we set the first Schur block to be 1 × 1 with real eig<u>en</u>value *λ*_1_. Since *U* is strictly upper triangular, *Ue*_1_ = 0 and hence *T e*_1_ = *λ*_1_*e*_1_. Using 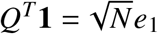, we obtain the row-sum identity:

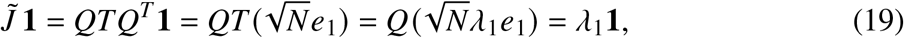

i.e., every row of 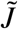 sums to *λ*_1_, which can be controlled directly via the spectrum (e.g., *λ*_1_ ≈ 0).

#### Assembly

Given sampled Λ, constructed *Q*, and LP solution *U*, we set 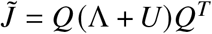 and recover 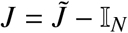.

### Amplifying initial conditions

Amplifying initial conditions are defined as perturbations that produce transient growth under the linearized dynamics. The most amplifying input direction *ϵ*_1_ is defined as the initial condition that maximizes the energy of the evoked neural response (Hennequin et al., 2014):

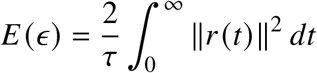

Because the system is stable, trajectories eventually return to baseline, ensuring that the response energy remains finite.

Following the procedure described in Hennequin et al. (2014), we construct a set of orthogonal input directions {*ϵ*_1_, *ϵ*_2_, …, *ϵ*_*N*_} ordered by their amplification strength. The leading direction *ϵ*_1_ corresponds to maximal amplification, while subsequent directions lead to progressively weaker transient responses.

### Transition inputs

To generate and switch between dynamical states, we apply brief transition inputs *I*_ext_(*t*) to the network dynamics (Eq. 1).

#### Initiating oscillatory or transient dynamics from baseline

Starting from baseline activity (*r* ≈ 0), brief inputs aligned with the leading amplifying direction of the Jacobian, *ϵ*_1_, initiate self-sustained oscillations. In contrast, inputs aligned with a less amplifying direction, for example *ϵ*_2_, elicit transient responses that decay back to baseline.

#### Resuming a periodic orbit from an arbitrary state

To drive the system toward a previously identified periodic orbit, we use a feedback controller that pulls the current state *r*(*t*) toward the orbit. At each time *t*, we find the nearest point *r*^*orb*^ (*t*) on the orbit and compute the error *e*(*t*) = *r*^orb^ (*t*) −*r*(*t*), and decompose it into components tangent and normal to the orbit. The resulting transition input is:

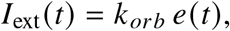

where *k*_*orb*_ sets how strongly the state is pulled toward the orbit.

#### Pulling activity toward a fixed point

To transition to a stable fixed point 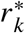 (including baseline or other steady states), we apply a similar feedback controller:

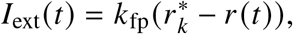

where *k*_fp_ sets how strongly the state is pulled toward the fixed point.

### Noise

#### Background noise

To assess robustness of the dynamics, we add temporally correlated noise to the system in the form of an Ornstein–Uhlenbeck (OU) process (Eq. 1). For each unit *i*, the noise variable *ξ*_*i*_ (*t*) has stationary variance

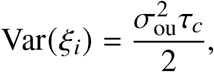

where *τ*_*c*_ = 10 ms is the noise correlation time and *σ*_ou_ is the diffusion coefficient.

Noise levels are chosen such that the average fluctuation amplitude at baseline corresponds to between 0.2% and 1.5% of the system’s dynamical range (Hennequin et al., 2014).

To compare robustness across networks with different amplitude of dynamics, the noise amplitude is normalized relative to the deterministic driving force. For a reference deterministic trajectory, we compute the root-mean-square (RMS) magnitude of the current 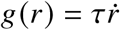.

The diffusion coefficient *σ*_ou_ is chosen such that the stationary per-coordinate standard deviation of the noise satisfies: std(*ξ*_*i*_) = *η*_dyn_ *g*_RMS_, where *η*_dyn_ is the target fraction of the deterministic drive. This yields

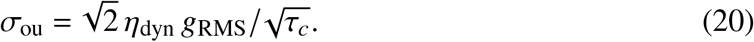

#### Initial-condition (input) noise

To assess sensitivity to initial condition perturbations, we add Gaussian noise to the initial condition:

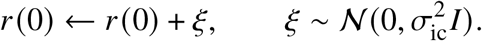

The expected perturbation norm scales with the dimensionality *d* of the system: 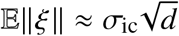. The per-coordinate standard deviation *σ*_ic_ is chosen such that E∥*ξ*∥ = *η*_ic_∥r(0) ∥, where *η*_ic_ is the target fraction of the unperturbed initial condition. Since the initial condition is normalized to ∥r(0) ∥ = 1, this yields

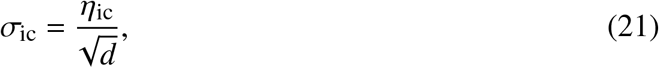

#### Critical noise amplitude

We define a deterministic reference trajectory segment and compare noisy trajectories against it using a tube criterion in full state space.

An ellipsoid is fit to the reference trajectory to determine a characteristic orbit size *S*. A tolerance tube of radius *ε* = *αS* with *α* = 0.1 is defined. A noisy trajectory is classified as remaining on the same orbit if at least 95% of its points lie within distance *ε* of the reference trajectory, and the maximum deviation does not exceed 2*ε*.

For each noise amplitude *η*, we run 100 stochastic trials. The critical noise amplitude is defined as the largest *η* for which at least 80 trials satisfy the tube criterion.

### Pseudospectra and stability

The *ε*-pseudospectrum of a matrix *J* is defined as in Trefethen and Embree, 2005:

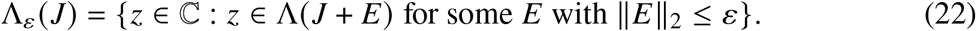

The *ε*-pseudospectrum consists of all complex numbers that become eigenvalues of *J* under perturbations of size at most *ε* in operator norm. An equivalent definition is

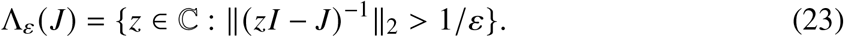

Thus, the pseudospectrum identifies regions in the complex plane where the resolvent norm is large, indicating high sensitivity of the spectrum to perturbations.

Here *J* denotes the Jacobian of the linearized system. The perturbation margin of *J* is defined as

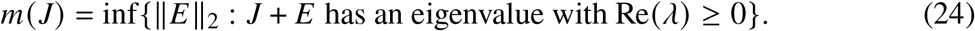

This quantity measures the smallest operator-norm perturbation required to move an eigenvalue to the imaginary axis and destabilize the system.

To compute the perturbation margin, we restrict *z* to the imaginary axis *z* = *iω*, which forms the boundary of the stability region. Using the resolvent definition,

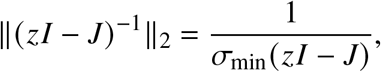

where *σ*_min_ denotes the smallest singular value. Hence,

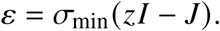

We evaluate *σ*_min_(*iωI* − *J*) across a range of frequencies *ω* and take the minimum value. This minimum equals the perturbation margin *m*(*J*).

### Gain modulation and frequency control of periodic orbits

Gain modulation is implemented as a multiplicative scaling of the incoming synapses of each neuron (Stroud et al., 2018):

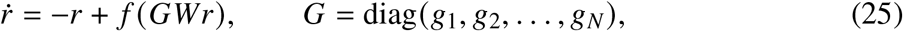

where *r* ∈ ℝ^*N*^ is the firing-rate vector, *W* ∈ ℝ^*N*×*N*^ is the recurrent connectivity matrix, *f* is applied elementwise, and *G* is a diagonal matrix of neuronal gains acting on postsynaptic excitatory neurons.

Let *F* (*g*) be the frequency of the stable limit cycle of the system as a function of the gain vector

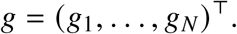

Our goal is to systematically control the oscillation frequency by adjusting *g*.

#### One-dimensional control direction in gain space

We parameterize the gain vector along a single direction in gain space:

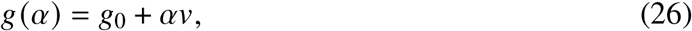

where *g*_0_ = (1, 1, …, 1)^⊤^ is the baseline gain vector, *v* ∈ ℝ^*N*^ is a direction in gain space, and *α* ∈ ℝ is a scalar control parameter.

Our objective is to identify a direction *v* such that varying *α* produces a locally monotonic and maximally sensitive change in frequency. This reduces frequency control to a single scalar “knob” *α*.

Formally, we seek *v* that maximizes the directional derivative of *F* at *g*_0_. By the chain rule,

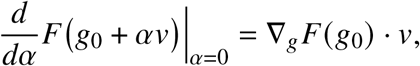

where ∇_*g*_ *F* (*g*_0_) is the gradient of the frequency function with respect to the gains.

This directional derivative is maximized when *v* is aligned with the gradient. Therefore, the direction of steepest frequency modulation is

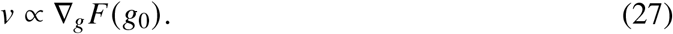

#### Numerical approximation of the gradient of *F*(*g*)

An analytical expression for the frequency function *F*(*g*) is not available. For a fixed gain vector *g*, we simulate the system until convergence to its stable limit cycle and estimate the frequency empirically.

We numerically approximate the gradient ∇_*g*_ *F*(*g*_0_) using finite differences. We apply a small perturbation *ε* to each gain *g*_*i*_, measure the resulting change in frequency and estimate the partial derivatives from these perturbations. Collecting these partial derivatives yields an estimate of the gradient vector, which defines the direction of steepest local frequency modulation.

#### Smooth frequency control

Under standard ODE theory, the vector field 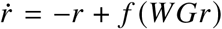 depends smoothly on the gain parameters *g*, provided that *f* is smooth. Consequently, as long as the limit cycle is not near a bifurcation point, small smooth perturbations of *g* induce small smooth changes in the orbit and in its associated period.

Therefore, for small |*α*|, the family of gain vectors *g*(*α*) generates a smooth family of limit cycles whose frequency varies smoothly with *α*. This provides a principled one-dimensional control direction in gain space that enables modulation of oscillation frequency.

### Bias-induced fixed points

We consider firing-rate dynamics with added constant bias

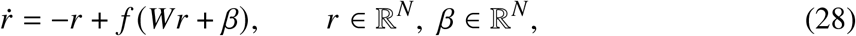

where the nonlinearity *f* is the shifted sigmoid (see Network Model) with fixed parameters *g* = 1, *θ* = 0, *r*_*max*_ = 1:

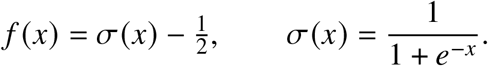

Useful identities of the nonlinearity are”

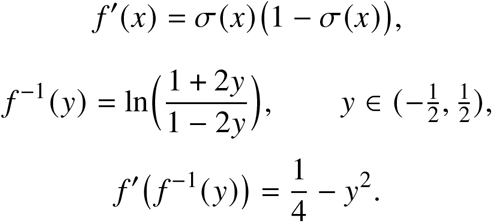

#### Reverse engineering the bias and Jacobian for a given fixed point

A fixed point *r*^∗^ satisfies:

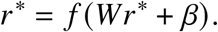

We want to investigate the nature of the emerging fixed points as we vary *β*. However, there is no straightforward mapping *β* ↦ *r*^∗^. Instead, applying *f* ^−1^ yields the equivalent relation

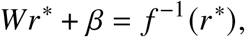

which implies that any chosen fixed point *r*^∗^ uniquely determines the bias:

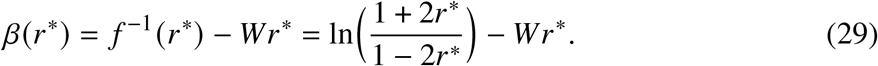

Furthermore, linearising the dynamics around the fixed point *r*^∗^ we obtain the Jacobian:

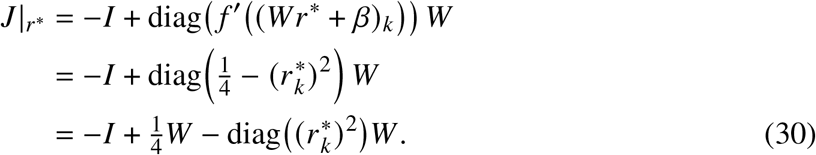

Local stability is determined by the spectral abscissa max_*i*_ ℜ*λ*_*i*_ *J* |_*r*_∗

#### Baseline dynamics (*β* = 0)

For zero bias *β* = 0, the fixed point of the system 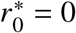 has corresponding Jacobian:

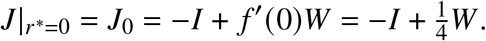

#### Uniform bias (*β* = *b*1)

Assuming a row-sum condition for the connectivity matrix *W* **1** = *s***1**, a uniform bias *β* = *b***1** admits a homogeneous fixed point: *r*^∗^ = *u*:

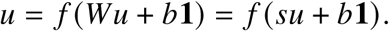

The Jacobian at the homogeneous fixed point is:

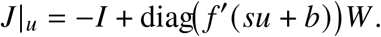

Since the slope *f*′(*su* + *b*) is constant across units and satisfies 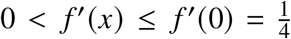, and because *J*_0_ = −*I* + *f*′(0)*W* is stable by construction, the Jacobian along the homogeneous branch is contracting as |*u*| increases. As the fixed point moves away from the origin, the slope *f*′(*su* + *b*) decreases monotonically, diminishing the recurrent contribution. In the limit of large bias magnitude, the Jacobian approaches *J* |_*u*_ → −*I*, corresponding to the purely contracting linear system 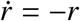.

#### Heterogeneous biases create “islands” of stable fixed points

For heterogeneous *β* — and hence a heterogeneous fixed point *r*^∗^ — slopes vary across units and the Jacobian becomes:

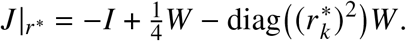

which no longer reduces to a scalar multiplication of *W*. Therefore, baseline stability of *J*_0_ does not imply stability of arbitrary heterogeneous fixed points, and stable solutions need not form a continuous branch. Stable heterogeneous fixed points are instead identified numerically.

#### Interpolation between fixed points

Given the baseline fixed point 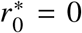 at *β*_0_ = 0 and a discovered stable heterogeneous fixed point 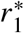 at bias *β*_1_, we define:

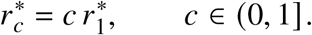

For each *c*, we can compute the bias that makes 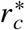 a fixed point:

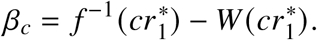

The Jacobian along this path is:

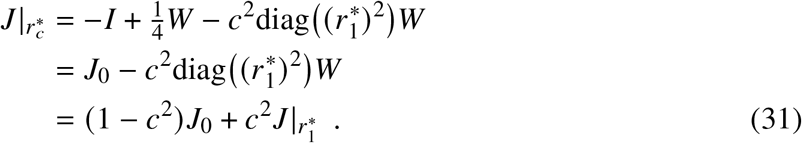

We analyse stability along this scaling path via 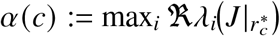.

#### Stability of homogeneous and heterogeneous fixed point paths

We compare the stability of homogeneous and heterogeneous fixed points at matched distance from baseline. We first numerically identify a stable heterogeneous fixed point 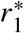 and compute its Euclidean distance from the baseline fixed point 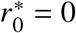:

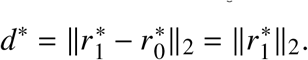

To construct a homogeneous fixed point at the same distance, we parameterize the homogeneous branch as 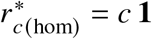. The coefficient *c* is chosen such that:

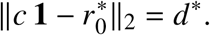

Since 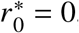, this reduces to:

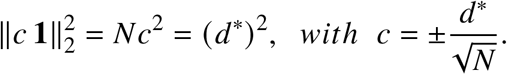

We select the solution satisfying 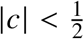 ensuring that 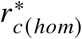 lies within the admissible range of the nonlinearity and *f* ^−1^ (*c*) is defined.

#### Separation between periodic orbits

When bias-induced stable fixed points support the existence of periodic orbits, we quantify how distinct the newly emerged orbits are in the full N-dimensional state space using a scale-normalized nearest-neighbor distance. For each orbit, we fit an ellipsoid from the principal axes of its covariance and scale it to match the orbit’s dimensions; the ellipsoid defines the characteristic orbit size *S*. Each orbit is then normalised by dividing by its own *S*.

To measure separation, we compute a directed nearest-neighbor distance. For each ordered pair of orbits (*A, B*), we take every point on *A*, find its nearest point on *B*, and record the 95th percentile of these point-wise distances. This yields a directed distance *d*_*A*ß*B*_. We take the larger of the two directions, *d*(*A, B*) = *max*(*d*_*A*ß*B*_, *d*_*B*ß*A*_). The resulting separation score is robust to outliers, respects the geometry of the trajectories, and reports how well two periodic orbits are separated relative to their intrinsic size.

#### Linear readouts

To decode low-dimensional movement trajectories *y*(*t*) from neural activity, we fit a single linear map from the population state *r*(*t*) ∈ ℝ^*N*^ to a two-dimensional output *y*(*t*) ∈ ℝ^2^ by weighted least squares. Training data are constructed from the deterministic periodic orbit trajectories stored for each state together with multiple noisy rollouts of those trajectories. Each orbit is paired with a target two-dimensional shape sampled at the orbit’s length. We additionally augment training with short noisy rollouts around stable fixed points, assigning them constant two-dimensional targets (e.g., a rest point).

### Repertoire dimensionality

We define the dynamical repertoire of a network as the collection of trajectories generated across bias-induced contexts (Eq. 26). So that orbits and transients are equally represented, we sample 50 contexts and include the limit cycle of each, and 50 transient trajectories of duration between 0.25 and 2.5 seconds drawn at random across those same 50 contexts.

Each trajectory *r* ^(*i*)^ (*t*) ∈ ℝ^*N*^ is mean-centered in time and rescaled such that its standard deviation equals 1. Without this normalization, the pooled statistics would be dominated by whichever trajectories happen to be initialized at the largest amplitudes; rescaling places long oscillations and short transients on equal footing without having to tune individual initial conditions to produce responses of comparable size.

### PCA analysis

#### Repertoire

Following Amematsro et al., 2025, we compare the dimensionality of individual trajectories with that of the full repertoire. We stack the normalized trajectories row-wise into 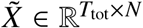, remove the per-neuron mean, apply PCA and define the repertoire dimensionality as the number of components required to explain 90% of the variance.

#### Subsets

We apply the same procedure to three additional sets of trajectories drawn from the network. These are the 50 orbits alone, the 50 transients alone, and a within-context set of one orbit and 49 transients drawn from a single bias-induced context. Reported dimensionalities are the number of components required to explain 90% of the variance of each subset.

#### Individual trajectories

We apply the same procedure to each normalized trajectory in isolation. The dimensionality of an individual trajectory is the number of components required to explain 90% of its variance.

#### Subspace overlap

To quantify how much two trajectories share their occupied subspace, we compute their pairwise subspace overlap as in Elsayed et al., 2016. For each normalized trajectory we compute its leading three principal components, defining a 3-dimensional subspace. The overlap from trajectory *i* to trajectory *j* is the variance of trajectory *j* captured by trajectory *i*’s subspace, divided by the variance *j* captures in its own top-3 subspace. Diagonal entries equal 1 by construction; off-diagonal entries report what fraction of one trajectory’s leading variance lies within another’s principal subspace.

## References

1 M. M. Churchland, J. P. Cunningham, M. T. Kaufman, J. D. Foster, P. Nuyu-jukian, S. I. Ryu, and K. V. Shenoy, “Neural population dynamics during reaching,” Nature 487 (2012).

2 K. V. Shenoy, M. Sahani, and M. M. Churchland, “Cortical control of arm movements: A dynamical systems perspective,” Annual Review of Neuroscience (2013).

3 M. T. Kaufman, M. M. Churchland, S. I. Ryu, and K. V. Shenoy, “Cortical activity in the null space: permitting preparation without movement,” Nature Neuroscience (2014).

4 G. F. Elsayed, A. H. Hara, M. T. Kaufman, M. M. Churchland, and J. P. Cunningham, “Reorganization between preparatory and movement population responses in motor cortex,” Nature Communications (2016).

5 A. H. Lara, J. P. Cunningham, and M. M. Churchland, “Different population dynamics in the supplementary motor area and motor cortex during reaching,” Nature Communications (2018).

6 A. A. Russo, S. R. Bittner, S. M. Perkins, J. S. Seely, B. M. London, A. H. Lara, A. Miri, N. J. Marshall, A. Kohn, T. M. Jessell, L. F. Abbott, J. P. Cunningham, and M. M. Churchland, “Motor cortex embeds muscle-like commands in an untangled population response,” Neuron 97 (2018).

7 S. Vyas, M. D. Golub, D. Sussillo, and K. V. Shenoy, “Computation through neural population dynamics,” Annual Review of Neuroscience (2020).

8 S. Saxena, A. A. Russo, J. Cunningham, and M. M. Churchland, “Motor cortex activity across movement speeds is predicted by network-level strategies for generating muscle activity,” eLife (2022).

9 E. A. Amematsro, E. M. Trautmann, N. J. Marshall, L. F. Abbott, M. N. Shadlen, D. M. Wolpert, and M. M. Churchland, “Motor cortex flexibly deploys a high-dimensional repertoire of subskills,” bioRxiv (2025).

10 K. C. Ames, S. I. Ryu, and K. V. Shenoy, “Neural dynamics of reaching following incorrect or absent motor preparation,” Neuron (2014).

11 C. Pandarinath, V. Gilja, C. H. Blabe, P. Nuyujukian, A. A. Sarma, B. L. Sorice, E. N. Eskandar, L. R. Hochberg, J. M. Henderson, and K. V. Shenoy, “Neural population dynamics in human motor cortex during movements in people with als,” eLife (2015).

12 A. K. Suresh, J. M. Goodman, E. V. Okorokova, M. Kaufman, N. G. Hatsopoulos, and S. J. Bensmaia, “Neural population dynamics in motor cortex are different for reach and grasp,” eLife (2020).

13 D. Sussillo and L. F. Abbott, “Generating coherent patterns of activity from chaotic neural networks,” Neuron (2009).

14 V. Mante, D. Sussillo, K. V. Shenoy, and W. T. Newsome, “Context-dependent computation by recurrent dynamics in prefrontal cortex,” Nature (2013).

15 D. Sussillo and O. Barak, “Opening the black box: Low-dimensional dynamics in high-dimensional recurrent neural networks,” Neural Computation (2013).

16 M. D. Golub, P. T. Sadtler, E. R. Oby, K. M. Quick, S. I. Ryu, E. C. Tyler-Kabara, A. P. Batista, S. M. Chase, and B. M. Yu, “Learning by neural reassociation,” Nature Neuroscience (2018).

17 J. P. Stroud, M. A. Porter, G. Hennequin, and T. P. Vogels, “Motor primitives in space and time via targeted gain modulation in cortical networks,” Nature Neuroscience (2018).

18 N. Maheswaranathan, A. Williams, M. Golub, S. Ganguli, and D. Sussillo, “Universality and individuality in neural dynamics across large populations of recurrent networks,” NeurIPS (2019).

19 G. R. Yang, M. R. Joglekar, H. F. Song, W. T. Newsome, and X.-J. Wang, “Task representations in neural networks trained to perform many cognitive tasks,” Nature Neuroscience (2019).

20 A. Dubreuil, A. Valente, M. Beiran, F. Mastrogiuseppe, and S. Ostojic, “The role of population structure in computations through neural dynamics,” Nature Neuroscience (2022).

21 L. N. Driscoll, K. V. Shenoy, and D. Sussillo, “Flexible multitask computation in recurrent networks utilizes shared dynamical motifs,” Nature Neuroscience (2024).

22 A. Colins Rodriguez, R. Fuentes-Flores, and M. D. Humphries, “Primary and supplementary motor cortex implement parallel control solutions for rhythmic and discrete arm movements,” bioRxiv (2026).

23 C. van Vreeswijk and H. Sompolinsky, “Chaos in neuronal networks with balanced excitatory and inhibitory activity,” Science (1996).

24 N. Brunel, “Dynamics of sparsely connected networks of excitatory and inhibitory spiking neurons,” Journal of Computational Neuroscience (2000).

25 T. P. Vogels and L. F. Abbott, “Signal propagation and logic gating in networks of integrate-and-fire neurons,” Journal of Neuroscience (2005).

26 T. P. Vogels, K. Rajan, and L. F. Abbott, “Neural network dynamics,” Annual Review of Neuroscience (2005).

27 H. Sompolinsky, A. Crisanti, and H. J. Sommers, “Chaos in random neural networks,” Physical Review Letters (1988).

28 K. Rajan, L. F. Abbott, and H. Sompolinsky, “Stimulus-dependent suppression of chaos in recurrent neural networks,” Physical Review E (2010).

29 S. Ostojic, “Two types of asynchronous activity in networks of excitatory and inhibitory spiking neurons,” Nature Neuroscience (2014).

30 J. Kadmon and H. Sompolinsky, “Transition to chaos in random neuronal networks,” Physical Review X (2015).

31 B. K. Murphy and K. D. Miller, “Balanced amplification: A new mechanism of selective amplification of neural activity patterns,” Neuron 61 (2009).

32 G. Hennequin, T. P. Vogels, and W. Gerstner, “Optimal control of transient dynamics in balanced networks supports generation of complex movements,” Neuron 82 (2014).

33 G. Christodoulou, T. P. Vogels, and E. J. Agnes, “Regimes and mechanisms of transient amplification in abstract and biological neural networks,” PLOS Computational Biology 18 (2022).

34 G. Christodoulou and T. P. Vogels, “The eigenvalue value (in neuroscience),” OSF Preprints (2022).

35 L. N. Trefethen and M. Embree, Spectra and Pseudospectra: The behavior of nonnormal matrices and operators (Princeton University Press, 2005).

36 M. S. Goldman, “Memory without feedback in a neural network,” Neuron (2009).

37 F. Mastrogiuseppe and S. Ostojic, “Linking connectivity, dynamics, and computations in low-rank recurrent neural networks,” Neuron (2018).

38 G. Bondanelli and S. Ostojic, “Coding with transient trajectories in recurrent neural networks,” PLOS Computational Biology 16 (2020).

39 L. Logiaco, L. F. Abbott, and S. Escola, “Thalamic control of cortical dynamics in a model of flexible motor sequencing,” Cell Reports (2021).

40 J. A. Gallego, M. G. Perich, L. E. Miller, and S. A. Solla, “Neural manifolds for the control of movement,” Neuron (2017).

41 M. G. Perich, D. Narain, and J. A. Gallego, “A neural manifold view of the brain,” Nature Neuroscience (2025).

42 G. Hennequin, E. J. Agnes, and T. P. Vogels, “Inhibitory plasticity: Balance, control, and codependence,” Annual Review of Neuroscience (2017).

43 Y. Ahmadian and K. D. Miller, “What is the dynamical regime of cerebral cortex?” Neuron (2021).

44 E. Marder and J.-M. Goaillard, “Variability, compensation and homeostasis in neuron and network function,” Nature Reviews Neuroscience (2006).

45 E. Marder, “Variability, compensation, and modulation in neurons and circuits,” PNAS (2011).

46 M. Khona and I. R. Fiete, “Attractor and integrator networks in the brain,” Nature Reviews Neuroscience (2022).

47 E. Salinas and P. Thier, “Gain modulation: A major computational principle of the central nervous system,” Neuron (2000).

48 F. S. Chance, L. F. Abbott, and A. D. Reyes, “Gain modulation from back-ground synaptic input,” Neuron (2002).

49 K. Thurley, W. Senn, and H.-R. Lüscher, “Dopamine increases the gain of the input-output response of rat prefrontal pyramidal neurons,” Journal of Neurophysiology (2008).

50 E. Marder, “Neuromodulation of neuronal circuits: back to the future,” Neuron (2012).

51 A. Colins Rodriguez, M. G. Perich, L. E. Miller, and M. D. Humphries, “Motor cortex latent dynamics encode spatial and temporal arm movement parameters independently,” Journal of Neuroscience (2024).

52 K. L. Briggman and W. B. J. Kristan, “Multifunctional pattern-generating circuits,” Annual Review of Neuroscience (2008).

53 P. T. Sadtler, K. M. Quick, M. D. Golub, S. M. Chase, S. I. Ryu, E. C. Tyler-Kabara, B. M. Yu, and A. P. Batista, “Neural constraints on learning,” Nature (2014).

54 J. A. Gallego, M. G. Perich, S. N. Naufel, C. Ethier, S. A. Solla, and L. E. Miller, “Cortical population activity within a preserved neural manifold underlies multiple motor behaviors,” Nature Communications (2018).

55 O. Marschall, D. G. Clark, and A. Litwin-Kumar, “A theory of multi-task computation and task selection,” bioRxiv (2025).

56 M. V. Tsodyks, W. E. Skaggs, T. J. Sejnowski, and B. L. McNaughton, “Paradoxical effects of external modulation of inhibitory interneurons,” Journal of Neuroscience (1997).

57 H. Ozeki, I. M. Finn, E. S. Schaffer, K. D. Miller, and D. Ferster, “Inhibitory stabilization of the cortical network underlies visual surround suppression,” Neuron (2009).

58 Y. Ahmadian, D. B. Rubin, and K. D. Miller, “Analysis of the stabilized supralinear network,” Neural Computation (2013).

59 A. Sanzeni, B. Akitake, H. C. Goldbach, C. E. Leedy, N. Brunel, and M. H. Histed, “Inhibition stabilization is a widespread property of cortical networks,” eLife (2020).

60 G. G. Turrigiano and S. B. Nelson, “Homeostatic plasticity in the developing nervous system,” Nature Reviews Neuroscience (2004).

61 T. P. Vogels, H. Sprekeler, F. Zenke, C. Clopath, and W. Gerstner, “Inhibitory plasticity balances excitation and inhibition in sensory pathways and memory networks,” Science (2011).

62 K. B. Hengen, A. Torrado Pacheco, J. N. McGregor, S. D. Van Hooser, and G. G. Turrigiano, “Neuronal firing rate homeostasis is inhibited by sleep and promoted by wake,” Cell (2016).

63 N. J. Priebe and D. Ferster, “Inhibition, spike threshold, and stimulus selectivity in primary visual cortex,” Neuron (2008).

64 L. Cimeša, L. Ciric, and S. Ostojic, “Geometry of population activity in spiking networks with low-rank structure,” PLOS Computational Biology (2023).

65 O. Yizhar, L. E. Fenno, T. J. Davidson, M. Mogri, and K. Deisseroth, “Optogenetics in neural systems,” Neuron (2011).

66 A. M. Packer, L. E. Russell, H. W. P. Dalgleish, and M. Häusser, “Simultaneous all-optical manipulation and recording of neural circuit activity with cellular resolution in vivo,” Nature Methods (2015).

67 Z. Zhang, L. E. Russell, A. M. Packer, O. M. Gauld, and M. Häusser,“Closed-loop all-optical interrogation of neural circuits in vivo,” Nature Methods (2018).

68 J. H. Marshel, Y. S. Kim, T. A. Machado, S. Quirin, B. Benson, J. Kad-mon, C. Raja, A. Chibukhchyan, C. Ramakrishnan, M. Inoue, J. C. Shane, D. J. McKnight, S. Yoshizawa, H. E. Kato, S. Ganguli, and K. Deisseroth, “Cortical layer–specific critical dynamics triggering perception,” Science (2019).

69 K. Daie, K. Svoboda, and S. Druckmann, “Targeted photostimulation un-covers circuit motifs supporting short-term memory,” Nature Neuroscience (2021).

70 E. K. Miller and J. D. Cohen, “An integrative theory of prefrontal cortex function,” Annual Review of Neuroscience (2001).

71 A. A. Disney, C. Aoki, and M. J. Hawken, “Gain modulation by nicotine in macaque V1,” Neuron (2007).

72 A. A. Disney and M. J. Higley, “Diverse spatiotemporal scales of cholinergic signaling in the neocortex,” Journal of Neuroscience (2020).

73 R. M. Kelly and P. L. Strick, “Cerebellar loops with motor cortex and prefrontal cortex of a nonhuman primate,” Journal of Neuroscience (2003).

74 A. C. Bostan and P. L. Strick, “The basal ganglia and the cerebellum: nodes in an integrated network,” Nature Reviews Neuroscience (2018).

75 E. Todorov and M. I. Jordan, “Optimal feedback control as a theory of motor coordination,” Nature Neuroscience (2002).

76 M. Schimel, T.-C. Kao, and G. Hennequin, “When and why does motor preparation arise in recurrent neural network models of motor control?” eLife (2024).

77 J. S. Calderón-García, G. Costalunga, T. P. Vogels, and D. Vallentin, “Interplay between syllable duration and pitch during whistle matching in wild nightingales,” Current Biology (2026).

78 A. J. Peters, S. X. Chen, and T. Komiyama, “Emergence of reproducible spatiotemporal activity during motor learning,” Nature (2014).

79 A. J. Peters, H. Liu, and T. Komiyama, “Learning in the rodent motor cortex,” Annual Review of Neuroscience (2017).

80 L. Duncker, L. N. Driscoll, K. V. Shenoy, M. Sahani, and D. Sussillo, “Organizing recurrent network dynamics by task-computation to enable continual learning,” in Advances in Neural Information Processing Systems (2020).

81 M. Beiran, A. Dubreuil, A. Valente, F. Mastrogiuseppe, and S. Ostojic, “Shaping dynamics with multiple populations in low-rank recurrent networks,” Neural Computation (2021).

82 W. F. Podlaski, E. J. Agnes, and T. P. Vogels, “High capacity and dynamic accessibility in associative memory networks with context-dependent neuronal and synaptic gating,” Physical Review X (2025).

83 W. Maass, T. Natschläger, and H. Markram, “Real-time computing without stable states: A new framework for neural computation based on perturbations,” Neural Computation (2002).

