## Supplementary material for "Dale’s law, stability and nonlinearity are sufficient constraints to ignite transient and self-sustained neural dynamics"

### Emergence of periodic orbits

We use an energy argument to show (i) how non-normal transient amplification can push dynamics away from a stable fixed point along specific directions, and (ii) how a saturating nonlinearity enforces an outer boundary such that network activity cannot grow unbounded. While this argument alone does not guarantee the existence of a limit cycle in $N$-dimensional systems, it isolates the two forces that shape the dynamics: amplification, which drives activity away from baseline, and saturating nonlinearity, which contains it.

**Decomposition of dynamics into linear and nonlinear components.**

Consider the rate dynamics:

$$\dot{r} = -r + f(Wr), \tag{S1}$$

where $r \in \mathbb{R}^N$, $W \in \mathbb{R}^{N\times N}$, and $f(\cdot)$ is applied elementwise (wit the same parameters across all neurons). $f$ is the parameterized shifted sigmoid chosen so that 0 is a fixed point of the system (Methods). The Jacobian at the origin is:

$$J \;=\; \frac{\partial}{\partial r}\big[-r + f(Wr)\big]\bigg|_{r=0} \;=\; -\mathbb{I}_N + f'(0)\,W. \tag{S1}$$

We assume that 0 is a *stable* fixed point, i.e. all eigenvalues of $J$ have negative real part, and that $J$ is *non-normal and amplifying*, i.e., for *preferred* input directions, the linearized dynamics exhibit transient growth before returning to baseline (Methods).

We define the nonlinear remainder:

$$n(r) := f(Wr) - f'(0)\,Wr. \tag{S2}$$

Then the dynamics can be written as the exact decomposition:

$$\begin{aligned}\dot{r} &= -r + f(Wr)\\ &= \big(-\mathbb{I}_N + f'(0)\,W\big)r + \big(f(Wr) - f'(0)\,Wr\big)\\ &= Jr + n(r)\end{aligned} \tag{S3}$$

where $Jr$ is the linear component and $n(r)$ is the nonlinear component.

We analyze the growth and decay of the activity norm $\|r\|$ using the standard quadratic energy function following (Bondanelli and Ostojic, 2020):

$$V(r) := \frac{1}{2}\|r\|^2. \tag{S4}$$

This function satisfies $V(r) > 0$ for all $r \neq r^*$ and $V(r^*) = 0$ at the fixed point $r^* = 0$. Differentiating $V$ gives:

$$\dot{V}(r) = r^\top \dot{r} = r^\top J r + r^\top n(r). \tag{S5}$$

If $\dot{V}(r) > 0$, the activity norm $\|r\|$ is increasing, whereas if $\dot{V}(r) < 0$, it is decreasing.

**Small-amplitude regime: $\|r\| \ll 1$.**

We investigate, in turn, the linear $r^\top J r$ and nonlinear $r^\top n(r)$ contributions.

**Linear contribution $\mathbf{r^\top Jr}$:** We decompose $J$ into a symmetric and a skew-symmetric part.

$$J = J_S + J_K, \qquad J_S := \frac{1}{2}(J + J^\top), \qquad J_K := \frac{1}{2}(J - J^\top).$$

Since $J_K^\top = -J_K$, then:

$$r^\top J_K r = 0, \qquad \forall r \in \mathbb{R}^N.$$

Hence:

$$r^\top J r = r^\top J_S r.$$

Let $v$ be a unit eigenvector of $J_S$ corresponding to its largest eigenvalue $\lambda^S_{\max}$, and write $r = \|r\|v$. Then

$$r^\top J r \;=\; (\|r\|v)^\top J_S(\|r\|v) \;=\; \lambda^S_{\max}\,\|r\|^2.$$

Amplification is associated with a positive numerical abscissa, $\lambda^S_{\max} > 0$, even when $J$ is stable (all eigenvalues have negative real part), implying the existence of directions along which the activity norm $\|r\|$ instantaneously grows:

$$\exists\, r \neq 0 \;\text{ s.t. }\; r^\top J r > 0.$$

**Nonlinear contribution $\mathbf{r}^\top\mathbf{n}(\mathbf{r})$:** Because $f$ acts elementwise on $Wr$, we Taylor-expand each component of the input independently. For the $i$-th component:

$$f((Wr)_i) \;=\; f(0) + f'(0)\,(Wr)_i + \tfrac{1}{2}f''(0)\,(Wr)_i^2 + O\Big((Wr)_i^3\Big).$$

Stacking the components into vector form, the quadratic term assembles into the elementwise square $(Wr)^{\odot 2}$, with $i$-th entry $(Wr)_i^2$. Using $f(0) = 0$, the remainder takes the form:

$$n(r) \;=\; f(Wr) - f'(0)\,Wr \;=\; \tfrac{1}{2}\,f''(0)\,(Wr)^{\odot 2} + O\Big(\|Wr\|^3\Big).$$

To bound $\|n(r)\|$, two standard inequalities suffice. First, the elementwise square satisfies $\|v^{\odot 2}\| \leq \|v\|^2$ for any vector $v$ and second, $\|Wr\| \leq \|W\|\,\|r\|$. For sufficiently small $\|r\|$, the cubic term is negligible, and combining the inequalities yields:

$$\|n(r)\| \;\leq\; c\,\|W\|^2\,\|r\|^2,$$

where the constant $c = |f''(0)|/2$. Applying the Cauchy–Schwarz inequality then gives $|r^\top n(r)| \leq \|r\|\,\|n(r)\| \leq c\,\|W\|^2\,\|r\|^3$, or equivalently:

$$-c\,\|W\|^2\,\|r\|^3 \;\leq\; r^\top n(r) \;\leq\; c\,\|W\|^2\,\|r\|^3.$$

Combining the equality for the linear component with the lower bound on the nonlinear component, along the most amplifying direction $v$, yields:

$$\dot V(r) \;=\; r^\top J r + r^\top n(r) \;\geq\; \lambda^S_{\max}\,\|r\|^2 \;-\; c\,\|W\|^2\,\|r\|^3. \tag{S9}$$

For $\|r\| < \lambda^S_{\max}/(c\,\|W\|^2)$, the cubic term is dominated by the quadratic, and the sign of $\dot V$ is dictated by $r^\top J_S r$. In particular, when $J$ is amplifying ($\lambda^S_{\max} > 0$), $\dot V(r) > 0$ along the amplifying direction $v$.

**Large-amplitude regime:** $\|r\| \gg 1$**.**

$f$ saturates elementwise, thus there exists $c_{\text{sat}} > 0$ such that:

$$|f(x)| \leq c_{\text{sat}}, \qquad \forall x \in \mathbb{R}.$$

Then $\|f(Wr)\| \leq c_{\text{sat}}\sqrt{N}$, $\forall r \in \mathbb{R}^N$. Starting from:

$$\dot V(r) = r^\top(-r) + r^\top f(Wr) = -\|r\|^2 + r^\top f(Wr),$$

we bound the cross term by the Cauchy–Schwarz inequality:

$$|r^\top f(Wr)| \leq \|r\|\,\|f(Wr)\| \leq \|r\|\,c_{\text{sat}}\sqrt{N}.$$

Hence:

$$\dot{V}(r) \leq -\|r\|^2 + c_{\text{sat}}\sqrt{N}\,\|r\| = -\|r\|\big(\|r\| - c_{\text{sat}}\sqrt{N}\big). \tag{S6}$$

Therefore, for $\|r\| > c_{\text{sat}}\sqrt{N}$, $\dot{V}(r) < 0$ for all directions. This implies the existence of an outer boundary: trajectories cannot escape to infinity and eventually decay once $\|r\|$ is sufficiently large.

The emerging picture is that: (i) near the origin ($\|r\| \ll 1$), non-normal transient amplification produces growth of the activity norm $\|r\|$ in specific (amplifying) directions; while (ii) far from the origin ($\|r\| \gg 1$), the saturating nonlinearity ensures that activity cannot grow without bound and eventually decays. In stable, Dalean, amplifying, nonlinear systems, these two conditions are *necessary but not sufficient* for the emergence of periodic orbits. How the linear and nonlinear components interact at *intermediate* $\|r\|$ determines the fate of the dynamics.

### Dynamics with positive firing rates

We consider the firing rate dynamics $\dot{r} = -r + f(Wr)$, where the elementwise nonlinearity $f$ is a shifted sigmoid chosen such that the origin is a fixed point, $f(0) = 0$:

$$f(x) = \frac{r_{max}}{1 + e^{-g(x-\theta)}} - \frac{r_{max}}{1 + e^{g\theta}} = \varphi(x) - \varphi(0), \tag{S7}$$

where $\varphi(\cdot)$ is the parameterised sigmoid with gain $g$, threshold $\theta$, and maximum firing rate $r_{max}$ (all positive).

This formulation allows activity to fluctuate around the baseline $r = 0$. To instead obtain dynamics with an arbitrary (generally positive) baseline, we consider the unshifted system with a bias vector $\beta$:

$$\dot{r} = -r + \varphi(Wr + \beta). \tag{S8}$$

Let $s := \varphi(0)$, and choose the bias $\beta = -Ws$. Then $r = s$ is a fixed point, since:

$$\varphi(Ws + \beta) = \varphi(0) = s.$$

Defining the translated coordinates:

$$y := r - s, \qquad r = y + s, \tag{S9}$$

and noting that $\dot{r} = \dot{y}$, we obtain:

$$\begin{aligned} \dot{y} &= -(y + s) + \varphi\big(W(y + s) + \beta\big) \\ &= -y - s + \varphi\big(Wy + (Ws + \beta)\big). \end{aligned} \tag{S10}$$

With $\beta = -Ws$, the constant input cancels, yielding:

$$\begin{aligned} \dot{y} &= -y - s + \varphi(Wy) \\ &= -y + \big(\varphi(Wy) - \varphi(0)\big) \\ &= -y + f(Wy). \end{aligned}$$

Therefore, the biased positive-rate system:

$$\dot{r} = -r + \varphi(Wr - Ws)$$

corresponds to a translation of the shifted system:

$$\dot{y} = -y + f(Wy),$$

where $r(t) = y(t) + s$.

Consequently, all trajectories and attractors are preserved up to a translation. Choosing a sigmoid satisfying $\varphi(0) > 0$ produces a strictly positive baseline. The repertoire of dynamics can be shifted to be positive by selecting $s$ sufficiently large to compensate for the negative excursions of $y(t)$.

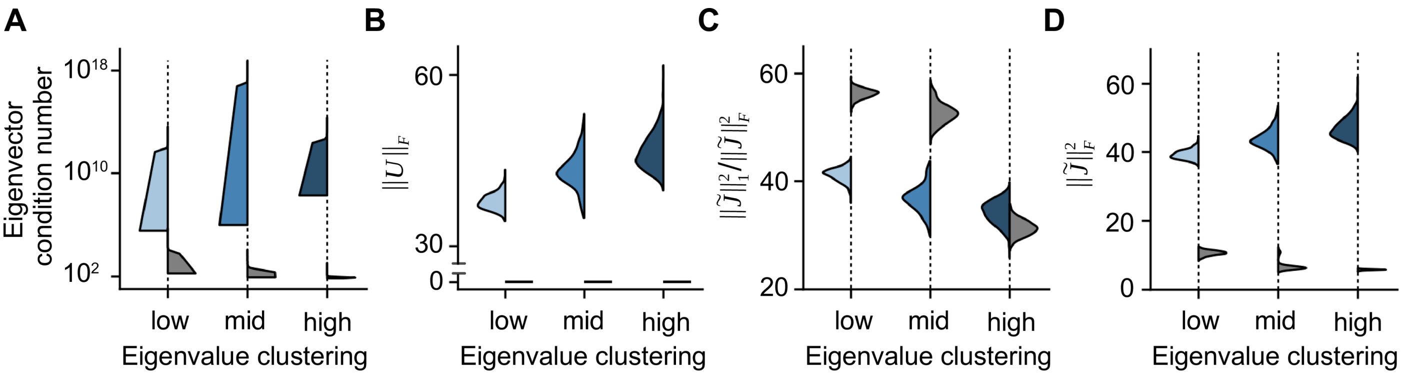


**Figure S1 | Mechanisms linking eigenvalue clustering to transient amplification.** **A)** Eigenvector condition number as a function of eigenvalue clustering for non-normal matrices with fixed strictly upper-triangular norm $U$ (blue) and normal controls (gray). Increasing clustering leads to larger condition numbers in non-normal matrices, reflecting increased eigenvector alignment. Normal matrices show the opposite trend as the spectrum collapses. **B)** Strictly upper-triangular norm $U$ as a function of eigenvalue clustering when the total Frobenius norm $\tilde{J}_F$ is held approximately fixed. As clustering increases, spectral variance decreases and energy shifts from the diagonal eigenvalue component into U, increasing $U$. Normal matrices have $U = 0$ by construction. **C)** Ratio $\tilde{J}_1^2/\tilde{J}_F^2$ under fixed $L1$ weight budget. Stronger clustering leads to a reduced $L1/L2$ ratio, indicating more concentrated weight mass. This effect is observed for both non-normal (blue) and normal (gray) matrices. **D)** Frobenius norm $\tilde{J}_F$ under fixed $L1$ budget. Increasing clustering leads to larger $\tilde{J}_F$, reflecting the emergence of fewer, larger-magnitude weights required to satisfy spectral constraints. Combined with panel b, this implies a further increase in the strictly upper-triangular component $U$.

### Clustering of eigenvalues increases transient amplification

We investigate how the distribution of eigenvalues of a stable Jacobian $J$ (all eigenvalues satisfy $\mathrm{Re}(\lambda_i) < 0$) influences the amount of transient amplification (see Methods) the system can generate. To this end, we use the shifted Jacobian $\tilde{J} = J + \mathbb{I}_N$, whose real Schur decomposition (see Methods) is:

$$\tilde{J} = QTQ^T, \qquad T = \Lambda + U, \qquad QQ^T = \mathbb{I}_N. \tag{S11}$$

$\Lambda$ is block-diagonal (the eigenvalues of $\tilde{J}$) and $U$ is strictly upper triangular. The matrix $U$ captures the non-normal structure responsible for amplification. We identify three complementary mechanisms through which eigenvalue clustering—the degree to which the eigenvalues are grouped together in the complex plane—increases amplification.

#### Clustering increases eigenvector overlap

Even when the strictly upper-triangular norm $\|U\|_F$ is fixed, clustering eigenvalues forces eigenvectors to align (Christodoulou et al., 2022). Consider the simple $2 \times 2$ upper-triangular case:

$$T = \begin{pmatrix} \lambda_1 & t \\ 0 & \lambda_2 \end{pmatrix}, \qquad t \in \mathbb{R}.$$

Eigenvectors satisfy:

$$v_1 = \begin{pmatrix} 1 \\ 0 \end{pmatrix}, \qquad v_2 = \begin{pmatrix} 1 \\ (\lambda_2 - \lambda_1)/t \end{pmatrix}.$$

As $\lambda_2 - \lambda_1 \to 0$, $v_2 \to v_1$, thus increasing eigenvector overlap. Under strong clustering, the eigenvector matrix becomes ill-conditioned, making the computation of the eigenvectors and of their overlap numerically unstable. Instead, we use the condition number of the eigenvector matrix as a proxy for overlap (Fig. S1A).

#### Clustering increases the strictly upper-triangular norm

The Frobenius norm is unitary invariant:

$$||\tilde{J}||_F^2 = ||T||_F^2.$$

Since $T = \Lambda + U$,

$$||\tilde{J}||_F^2 = \sum_{i=1}^{N} |\lambda_i|^2 + ||U||_F^2,$$

and therefore

$$||U||_F^2 = ||\tilde{J}||_F^2 - \sum_{i=1}^{N} |\lambda_i|^2.$$

We increase eigenvalue clustering while keeping the eigenvalue mean $\mu$ approximately fixed:

$$\sum_{i=1}^{N} |\lambda_i|^2 = N|\mu|^2 + \sum_{i=1}^{N} |\lambda_i - \mu|^2.$$

The second term, $\sum_{i=1}^{N} |\lambda_i - \mu|^2$, is N times the variance of the eigenvalues. Increasing clustering decreases the variance, and hence decreases $\sum_i |\lambda_i|^2$ as well. For approximately fixed $||\tilde{J}||_F^2$, this forces $||U||_F^2$ to increase. Thus, clustering shifts matrix "energy" away from the diagonal eigenvalue component $\Lambda$ and into the strictly upper-triangular non-normal component $U$, directly enhancing transient amplification (Fig. S1B).

#### Clustering concentrates synaptic weights into fewer, stronger connections

Under a fixed synaptic weight budget — the $L^1$ norm $||\tilde{J}||_1$ — tighter clustering requires concentrating weight into fewer, larger entries. As a result, the $L^1/L^2$ ratio $||\tilde{J}||_1^2/||\tilde{J}||_F^2$ decreases (Fig. S1C) and the Frobenius norm $||\tilde{J}||_F$ grows (Fig. S1D). A larger $||\tilde{J}||_F$ adds to the previous mechanism: since clustering simultaneously lowers $\sum_i |\lambda_i|^2$, the non-normal component $||U||_F^2 = ||\tilde{J}||_F^2 - \sum_{i=1}^{N} |\lambda_i|^2$ grows on both counts.

Overall, increasing eigenvalue clustering enhances transient amplification through (i) increasing eigenvector overlap, (ii) increasing the norm of the strictly upper-triangular component $U$, and (iii) concentrating synaptic weights into fewer, stronger connections. Together, these effects increase amplification while the real parts of eigenvalues remain strictly negative and stability is preserved.

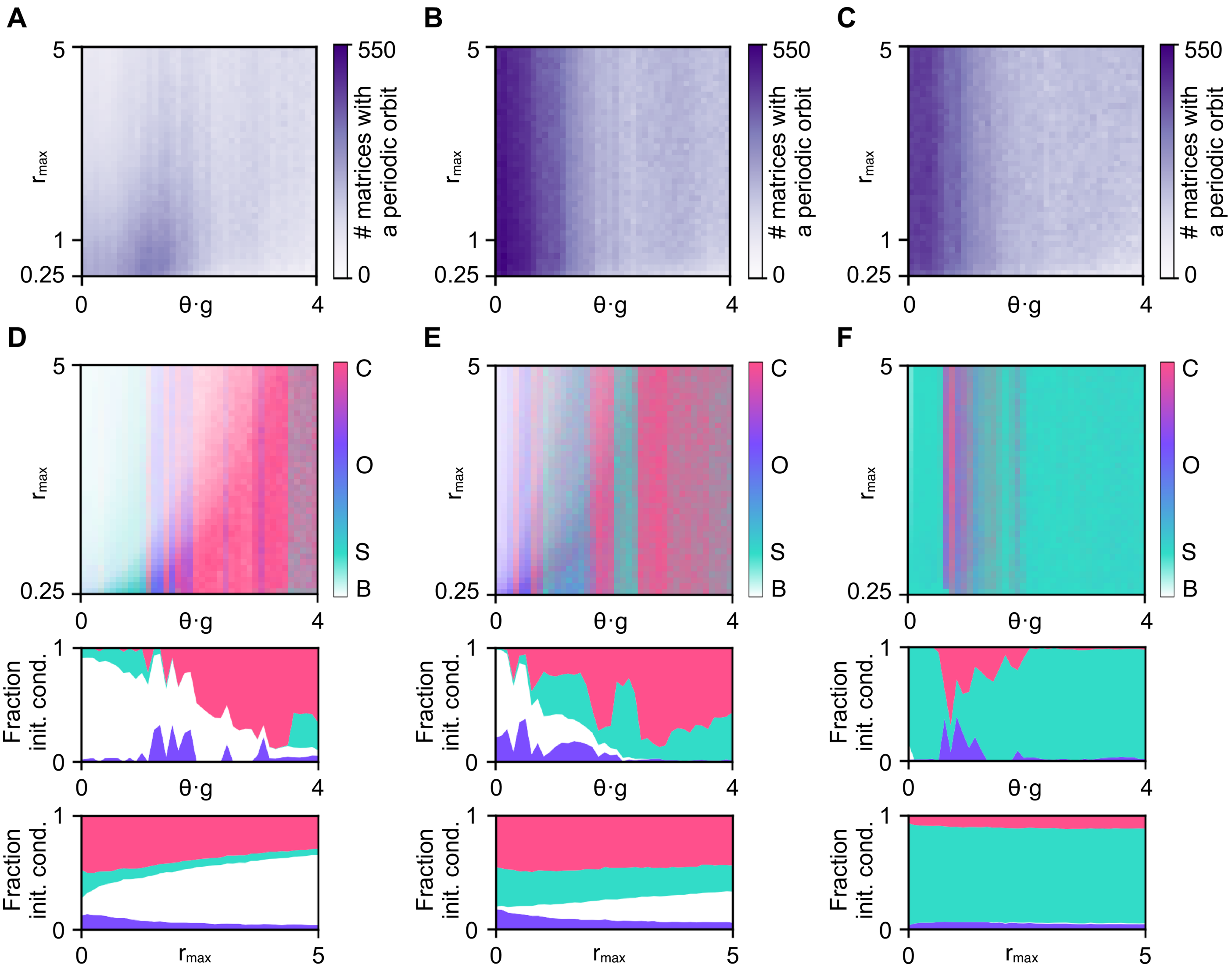


**Figure S2 | Nonlinearity selects the attractor.** **A–C** Number of matrices with at least one periodic orbit in the low (A), medium (B), and high (C) eigenvalue clustering ensembles, as a function of the nonlinearity parameters: the product of threshold and gain ($\theta \cdot g$), and range ($r_{max}$). **D–F** Dynamics of one example network from the low (D), medium (E), and high (F) clustering ensembles as a function of nonlinearity parameters (the networks are the same as in Fig. 4D-F). **Top:** for every combination of $\theta \cdot g$ and $r_{max}$, each network is initialized with 100 orthogonal initial conditions (Methods). Color indicates the fraction of trajectories converging to chaos (C, pink), a non-baseline steady state (S, teal), or a periodic orbit (O, purple); saturation indicates the fraction returning to baseline (B). **Middle:** fraction of initial conditions converging to each dynamical state (B, O, S, C) as a function of $\theta \cdot g$, averaged over $r_{max}$. **Bottom:** same, as a function of $r_{max}$, averaged over $\theta \cdot g$.
